# Mental replay rescales the subjective duration of past events via alpha and theta oscillations

**DOI:** 10.64898/2026.10.25.761606

**Authors:** Winny Wing Yin Yue, Zexuan Mu, Ziqing Yao, Danni Chen, Yuqi Zhang, Jing Liu, Xiaoqing Hu

**Author notes:** Corresponding Authors: Xiaoqing Hu, Department of Psychology, The University of Hong Kong, Jing Liu, School of Psychology, South China Normal University.

## Abstract

Human memory recall is flexible in time, allowing past events to be mentally replayed at variable speeds. However, it remains unclear whether temporally rescaled replay distorts the perceived duration of remembered events, and whether such distortion is tied to changes in the retrieved content. Here we instructed participants to mentally replay previously encoded naturalistic videos at compressed, expanded, or self-paced (free) speeds while recording scalp electroencephalography (EEG). Replaying an event at an expanded pace lengthened participants’ subsequent judgments of its original duration relative to free and compressed replay, and the longer an event had been replayed, the longer it was judged to have lasted. Crucially, recalled memory content was unaffected, with gist accuracy and recalled detail comparable across speed conditions. During replay, alpha and theta power were reduced under compression, and both interacted with the speed instruction in predicting reported duration. Specifically, alpha power exhibited a stronger positive relationship with duration judgments under compressed replay than under free or expanded replay. Conversely, the relationship between theta power and reported duration was significantly more positive under expanded replay than under free or compressed replay. Neural pattern similarity between instructed and later spontaneous replay varied with replay speed, yet this reinstatement neither predicted reported duration nor mediated the oscillatory effects. These findings suggest that remembered duration is not read out from the amount of content retrieved but reconstructed at retrieval through oscillatory dynamics, decoupling the subjective scaling of time from the content of memory.

## Introduction

A defining feature of episodic memory is its temporal flexibility: the human brain can compress hours-long experience into a snapshot or, conversely, expand a fleeting moment into a detailed prolonged replay (Bellingrath, 2023; Jeunehomme et al., 2020; Jeunehomme & D’Argembeau, 2019; Leroy et al., 2024; Michelmann et al., 2019). Notably, altering the speed of post-encoding mental replay allows us to efficiently navigate the past, engage in prospective planning, and make adaptive decisions (Biderman et al., 2020; Howard, 2018; Schacter et al., 2007). However, it remains unclear whether this temporally rescaled memory replay reshapes one’s existing perception of an event’s duration and, if so, whether it correspondingly alters the retrieved content.

The prevailing “storage size” hypothesis posits that retrospective judgments or estimates of event durations are inferred based on the amount or “richness” of the recalled event (Burt & Kemp, 1991; Faber & Gennari, 2015; Ornstein, 1969). Under this framework, subjective time is a byproduct of retrieved memory content; memories that are rich in perceptual and semantic details are often judged to be longer (D’Argembeau et al., 2022; Eagleman & Pariyadath, 2009; Jeunehomme et al., 2018; Jeunehomme & D’Argembeau, 2019). Moreover, increasing the number of event changes or segments consistently lengthens retrospective duration estimates (Block & Reed, 1978; Clewett et al., 2019; Faber & Gennari, 2015; Poynter, 1983; Yue et al., 2026; Zakay & Block, 2004). Converging clinical evidence also links the two: memory deficits, such as those associated with Alzheimer’s disease, are often accompanied by distortions in time perception (El Haj et al., 2013; El Haj & Kapogiannis, 2016a; Mioni et al., 2014). These findings suggest that duration judgement may scale with the richness of the retrieved memory content.

In contrast, recent neurophysiological evidence indicates an independent temporal storage account, which proposes that the brain represents temporal structure independently of sensory or semantic representations. In rodents, hippocampal “time cells” fire sequentially to bridge delays, encoding the passage of time even without changes in external input (Cao et al., 2022; Eichenbaum, 2014; MacDonald et al., 2011; Mau et al., 2018; Pastalkova et al., 2008). In humans, functional magnetic resonance imaging (fMRI) studies show that, for events separated by the same objective interval, greater hippocampal pattern similarity predicts that they are later remembered as closer in time, even when they span different contexts (Ezzyat & Davachi, 2014). This suggests that hippocampal activity carries a representation of temporal distance.

Crucially, studies with human single-unit recordings have identified distinct populations of time cells and ramping cells that map elapsed time irrespective of the surrounding environmental context or experiential content. Together, these findings suggest that duration judgement and memory content retrieval possibly rely on dissociable neural representations and neurocomputational processes (Reddy et al., 2021; Umbach et al., 2020).

Although not mutually exclusive, the two accounts diverge on whether remembered duration can change without a corresponding change in memory content, a prediction that is difficult to test because evidence for each has come from different levels of analysis (behavioural versus neural). Existing experimental paradigms typically manipulate event duration by altering the amount of information encoded, e.g., adding more segments or increasing stimulus complexity, thereby conflating the “time” variable with the “content” variable (Block, 1992; Clewett et al., 2019; Eagleman & Pariyadath, 2009; Matthews & Meck, 2016). Consequently, it remains unclear how actively manipulating the temporal pacing of a memory—such as altering its mental replay speed while holding content constant—impacts perceived duration, subsequent episodic retrieval, and its underlying neural mechanism.

For the underlying oscillatory mechanisms, we particularly focused on alpha and theta oscillations, which might serve complementary roles in timing and memory structuring. Specifically, alpha oscillatory activity (8–12 Hz) has long been proposed as a primary pacemaker for internal timing (Azizi et al., 2023; Klimesch, 2012; Kononowicz & van Wassenhove, 2016). Mechanistically, higher alpha power reflects active neuronal inhibition that gates information flow (Jensen & Mazaheri, 2010; Klimesch, 2012; Strauß et al., 2014) and rises with memory load during retention, indexing stronger suppression of task-irrelevant processing as demands increase (Jensen et al., 2002; Klimesch et al., 2007). This gating is dynamic: a “closed gate” (high alpha-band power) shields maintained content and resists updating, whereas an “open gate” (low alpha-band power) admits new information (Manza et al., 2014; Tu et al., 2025). During replay, the instructed pace may conflict with the duration encoded during viewing. We therefore hypothesized that alpha power regulates whether the instructed replay pace updates the encoded temporal structure of an event. Reduced alpha power should permit this updating, rendering reported duration sensitive to the instruction, whereas elevated alpha power should constrain it. Accordingly, alpha power should be positively associated with reported duration under compressed replay and negatively associated under expanded replay.

In contrast, theta activity organizes the sequential structure of memory. Transient hippocampal theta bouts align with the internal segmentation of experience, effectively establishing the discrete “time steps” of mental simulation (Herweg et al., 2020; Seeber et al., 2025). Furthermore, increased theta power reflects the binding of these discrete segments into a cohesive episodic sequence (Clewett et al., 2019; Fuentemilla et al., 2014; Heusser et al., 2016; Lisman & Jensen, 2013). With more and finer segments, this sequence is lengthened and takes longer to replay (Faber & Gennari, 2015; Jeunehomme & D’Argembeau, 2020). We therefore reasoned that expanded replay, as compared to compressed replay, might encourage an event to be re-segmented and re-experienced at a finer scale, lengthening the reported duration accordingly, with increased theta power supporting expanded temporal reconstruction and event binding.

To address these questions, we developed a novel “mental replay” paradigm to manipulate the post-encoding replay speed of memory while instructing participants to retain its full content. Participants viewed video episodes and were immediately instructed to mentally replay them at either Compressed, Expanded, or Free replay speed, before memory recall and duration judgments (Fig. 1). This paradigm allows us to determine how replaying event memories at different speeds would modulate neural oscillations and impact subsequent episodic memories, including the duration and content. By combining high-density electroencephalogram (EEG) with Representational Similarity Analysis (RSA), we demonstrate that retrospective temporal judgments operate independently of memory representation quality or content richness. Furthermore, we show that alpha and theta power differ with replay speed and track reported duration differently across conditions.

**Fig. 1.**
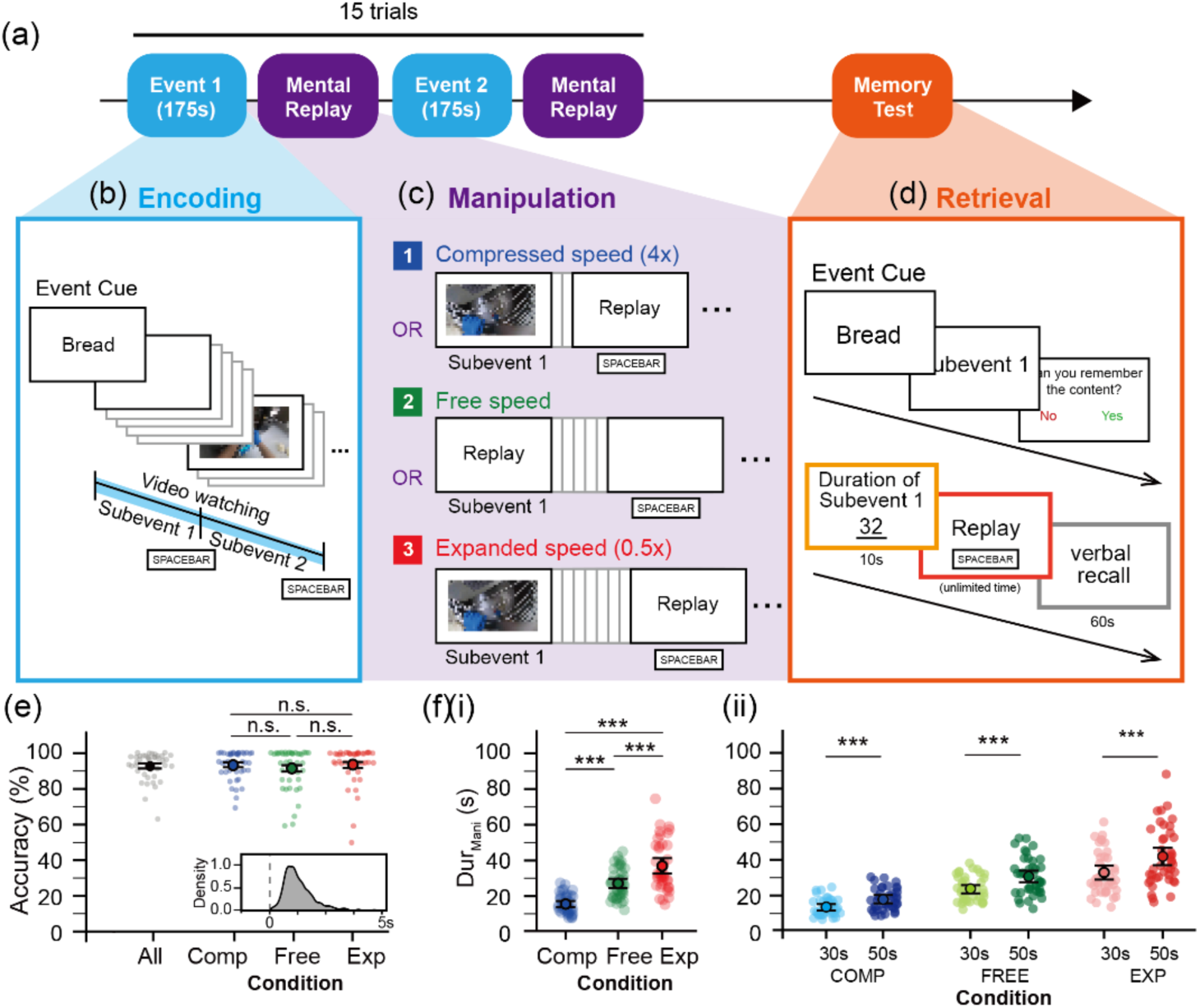
Experimental design and manipulation check results. (a) Overall experimental timeline. Participants completed 15 event trials, each consisting of video encoding (175 s) followed immediately by mental replay manipulation. A memory test was administered after all encoding and manipulation trials were completed. (b) Encoding phase. (c) Manipulation phase. Each video was randomly assigned to one of the three manipulation conditions, Compressed speed (Compress), Free speed (Free), Expanded speed (Expand), with 5 videos per condition. (d) Retrieval phase. (e) Boundary detection accuracy during encoding. Accuracy was calculated as the percentage of spacebar presses occurring within 5 seconds of actual subevent boundaries. Error bars represent standard error of the mean. (f)(i) Manipulation duration results. Mean replay duration during manipulation across conditions, confirming successful speed manipulation (Compress < Free < Expand). (ii) by original subevent duration (Duration_OriginalVideo_; 30 s vs. 50 s) within each condition. Individual transparent dots represent participant means; Solid color dots indicate group means with standard error. Compress: blue, Free: green, Expand: red. \*\*\**p* < .001, n.s. = not significant.

## Results

### Replay speed manipulation shapes duration judgments but not memory content

To test whether subjective duration can be reconstructed independently of memory content, participants encoded a series of naturalistic video events and then mentally replayed each video at an instructed speed (Compress, Free or Expand). In all conditions, participants were asked to replay the entire event in order without skipping any content, so that the instruction targeted the speed of retrieval rather than what was retrieved. In a subsequent test phase, we assessed for each subevent participants’ judgment of its original duration, the time they took to replay it spontaneously, and the gist and detail of their verbal recall (Fig. 1).

Each video comprised five subevents separated by clear event boundaries (Fig. 1b). During encoding, participants pressed a key at each subevent transition, confirming that they segmented the videos as intended and defining the duration of each subevent for later judgments. Boundary detection was highly accurate across all conditions (*M* = 93%, *SD* = 0.078; Fig. 1e). Moreover, no significant differences were found between Compress, Expand, and Free conditions for detection accuracy (Friedman test: *χ²(*2) = 1.197, *p* = .550) or response time (stats). At the neural level, boundary detection elicited expected EEG spectral power changes, including early delta-theta enhancement (∼1–8 Hz, 0–2.3s post-boundary) and subsequent alpha–beta suppression (∼8–30 Hz, 0.8–2.8s), consistent with prior reports of event boundary processing (Engel & Fries, 2010; Klimesch, 2012) (Supplementary Figure 1).

In the following mental replay phase, behavioural results confirmed that participants successfully implemented the mental replay speed instructions, with Compress showing the shortest manipulation replay duration (Duration_Mani_; *M* = 15.403 sec, *SD* = 8.147), followed by Free (*M* = 27.289 sec, *SD* = 11.927), and Expand (*M* = 38.231 sec, SD = 19.082), *H*(2) = 63.633, *p* < .001, with all pairwise comparisons significant (*ps* < .001) (Fig. 1f). Mental replay in the Free condition yielded a mean compression rate (mental replay duration divided by original subevent duration) of 0.703, consistent with spontaneous mental replay dynamics reported in previous studies (Jeunehomme et al., 2018; Jeunehomme & D’Argembeau, 2019) (Supplementary Figure 2). In addition, participants reliably tracked subevent length, replaying the short (30 s) subevents for less time than the long (50 s) subevents across all conditions (*χ²*(5) = 132.776, *p* < .001)(Supplementary Figure 3). These results provide converging evidence that participants successfully followed the mental replay speed instructions.

We next examined the impact of mental replay speed manipulation on subsequent temporal memory. Temporal memory was indexed in two ways: participants’ subjective estimation of the replayed event’s overall duration (Duration_TestReported_), and objective elapsed time participants spent actively replaying the video at test (Duration_TestReplay_). Although Duration_TestReported_ and Duration_TestReplay_ were significantly positively correlated (*r* = 0.482, *p* < .001) (Supplementary Figure 3), our primary analyses focused on the subjective estimation of the duration (Duration_TestReported_), as it directly indexes the subjective representation of time without being confounded by retrieval disfluency—such as hesitations or cognitive effort— that may inflate physical replay time (Duration_TestReplay_).

Employing a Generalized Linear Mixed Model (GLMM), which controlled for original subevent duration (30 vs. 50 s) and included random intercepts for participants and events, we found that speed manipulation directly and significantly influenced Duration_TestReported_ (*χ²*(2) = 17.889, *p* < .001, Fig 2a). Specifically, the Expand condition resulted in significantly longer reported durations compared to both the Free baseline (*b* = 0.055, *z* = 2.787, *p* = .005) and the Compress condition (*b* = 0.081, *z* = 4.155, *p* < .001), establishing a robust dilation of subjective time. In contrast, the difference between Compress and Free conditions did not reach significance (*p* = .182). Importantly, despite this dilation, participants preserved baseline accuracy by reliably judging 50-s events as longer than 30-s events (*b* = 0.21, *z* = 4.773, *p* < .001) (Fig. 2aii). Building on these categorical differences, a continuous measure of the manipulation replay duration (Duration_Mani_) strongly predicted Duration_TestReported_ (*χ²*(1) = 70.113, *p* < .001), with longer Duration_Mani_ associated with longer Duration_TestReported_ (*b* = 0.082, *z* = 8.373, *p* < .001) (Fig. 2aiii). By contrast, replay speed did not affect the time participants took to replay each subevent at test (Duration_TestReplay_; Supplementary Figure 4 and 5) These findings further suggest that subjective retrospective duration reports (Duration_TestReported_) provide a more sensitive behavioral metric for capturing subjective temporal scaling.

**Fig. 2.**
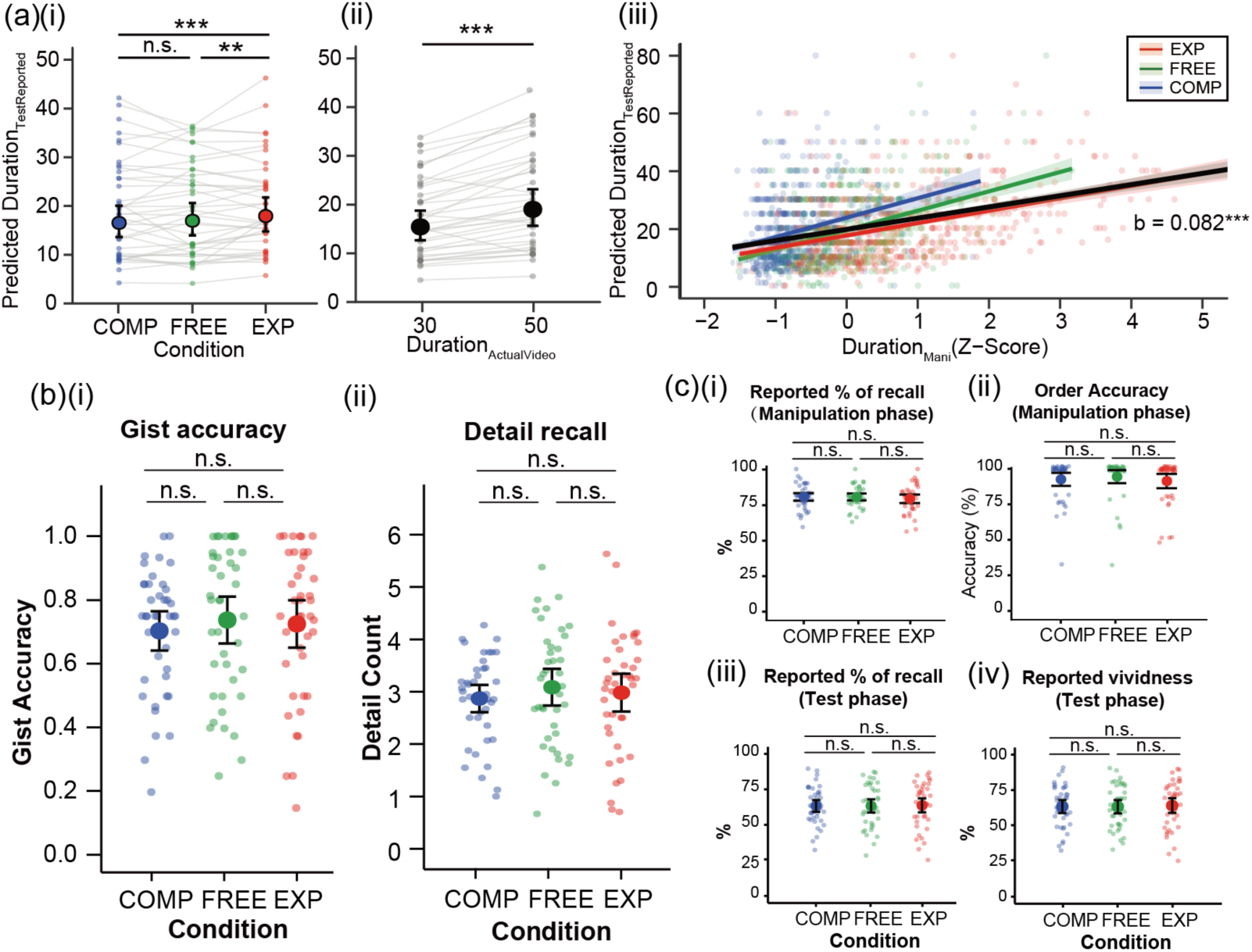
Speed manipulation alters temporal processing while preserving memory content. (a) Subjective duration estimates (Duration_TestReported_) during the test phase. (i) Duration_TestReported_ plotted across conditions. (ii) Duration_TestReported_ by Duration_ActualVideo_ (30s vs. 50s). (iii) Relationship between Duration_Mani_ (Z-scored) and subsequent Duration_TestReported_. The solid black line indicates the model fixed effect; colored lines represent condition-specific fits with shaded areas denoting 95% CI. (c) Memory recall of subevents across conditions. (i) Gist accuracy. (ii) Accurate detail recall count. (d) Additional memory and subjective measures across conditions during manipulation and test phases. (i) Self-reported percentage of recall during the manipulation phase. (ii) Temporal order accuracy during the manipulation phase. (iii) Self-reported percentage of recall during the test phase. (iv) Self-reported vividness of recall during the test phase. For all panels, large solid dots represent model-estimated marginal means or group means; error bars indicate 95% confidence intervals. Transparent dots represent individual participant means, connected by grey lines in (a) and (b). Compress: blue, Free: green, Expand: red. \*\**p* < .01, \*\*\**p* < .001, n.s. = not significant.

Crucially, the effect of replay speed on Duration_TestReported_ was not accompanied by changes in memory content. We analyzed participants’ verbal recall during the test phase, transcribing recordings and scoring gist accuracy and detail recall against pre-established templates (see Methods). Gist memory accuracy did not differ across conditions (Friedman test: *χ²*(2) = 2.545, *p* = .280, *BF_01_* = 8.18; Fig. 2bi). Similarly, detail recall — measured by the number of correctly remembered elements such as characters, actions, and objects — showed no significant condition effects (*χ²*(2) = 4.429, *p* = .109, *BF_01_* = 6.35; Fig. 2bii). No significant differences in self-reported recall percentages, recall vividness, and temporal order accuracy were observed across conditions (all *ps* > .206; Fig. 2c). Together, these results establish a clear behavioral dissociation: the speed manipulation reshaped the subjective feeling of time passage without influencing the richness of remembered content.

### Alpha and theta oscillatory power differently interact with replay speed in predicting duration judgments

We next examined how replay speed affected oscillatory activity during the manipulation phase. Because replay durations differed across conditions by design, power was estimated in 2-s sliding windows and averaged within each subevent, so that estimates did not scale with epoch length (see Methods). We used the Fitting Oscillations & One Over F (FOOOF) spectral parameterization to isolate the spectral oscillatory components from the fractal component (Fig. 3a; see Methods). Power differed significantly across replay speed for both alpha (*χ²*(2) = 55.629, *p* < .001) and theta (*χ²*(2) = 12.845, *p* = .002) (Fig. 3b). Alpha power was lower in Compress than in both Free (*b* = −0.031, *t* = −5.046, *p* < .001) and Expand (*b* = −0.044, *t* = −7.293, *p* < .001) conditions, while Free and Expand conditions did not differ (*b* = −0.013, *t* = −2.224, *p* = .067). Similarly, theta power was significantly lower in Compress than in both Free (*b* = −0.023, *t* = −3.112, *p* = .005) and Expand (*b* = −0.022, *t* = −3.113, *p* = .005), with no difference between Free and Expand (*b* = 0.000, *t* = 0.021, *p* = 1.000). Contrasts for the remaining bands are reported in Supplementary Figure 6a.

**Fig. 3.**
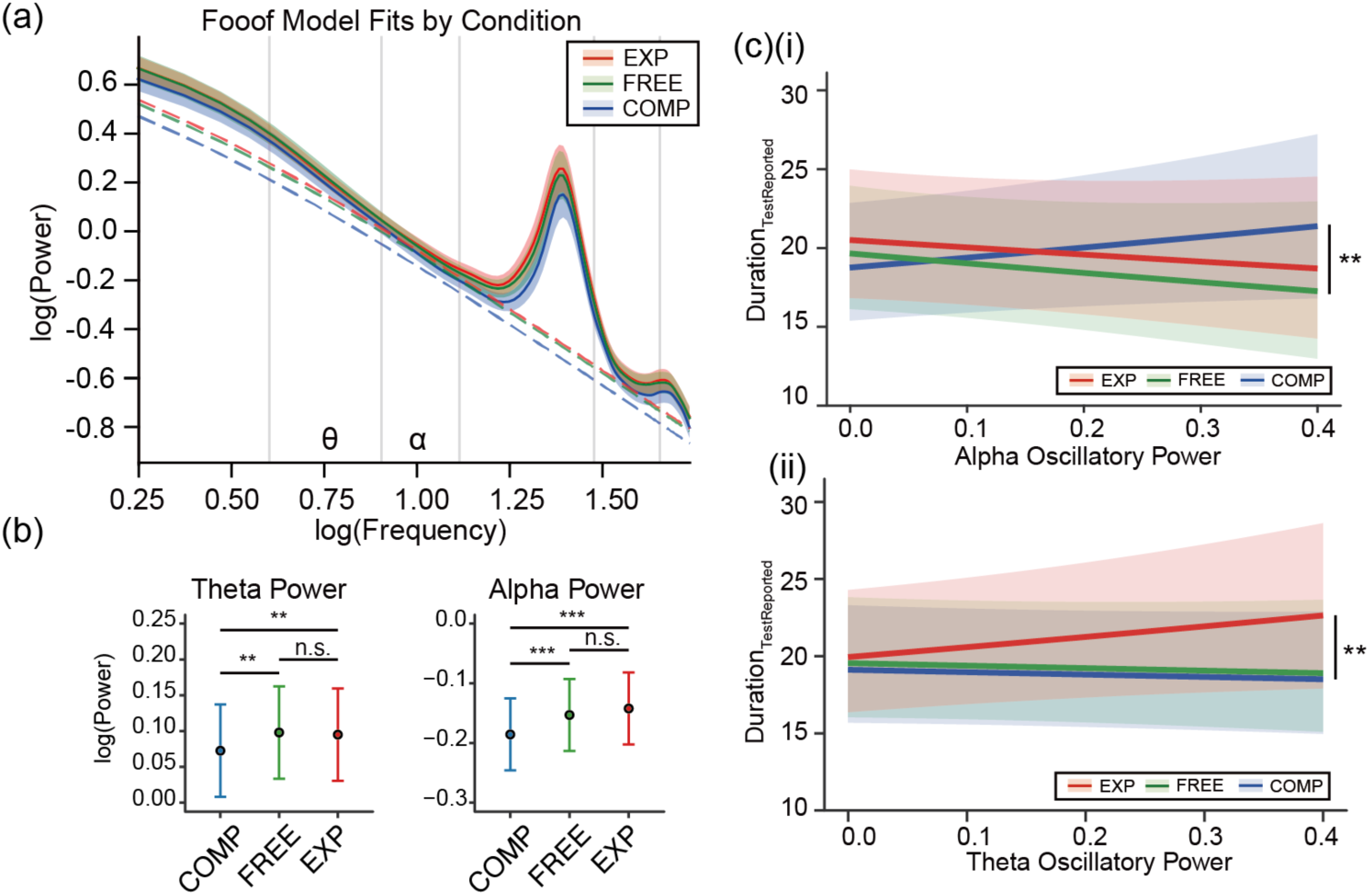
Distinct oscillatory mechanisms index subjective duration. (a) Neural power spectra parameterized via the FOOOF algorithm. The plot displays mean log-power across frequencies for each condition, with dashed lines representing the aperiodic fit. Shaded regions indicate SEM. Highlighted frequency bands are labelled along the x-axis (θ, α). (b) Estimated marginal means of log-transformed power across conditions for the theta and alpha frequency band. Error bars indicate 95% CI. (c)Dissociable oscillatory predictors of subjective duration judgement (Duration_TestReported_). (i) Predicted relationship between alpha oscillatory power and Duration_TestReported_; the Compress line exhibits a distinct positive slope relative to other conditions (indicated by the bracket). (ii) Predicted relationship between theta oscillatory power and Duration_TestReported_; the Expand line exhibits a distinct positive slope relative to other conditions. For all panels, condition color coding: Compress (blue), Free (green), Expand (red). Shaded ribbons represent 95% CI. \**p* < .05, \*\**p* < .01, \*\*\**p* < .001, n.s. = not significant.

To determine how neural oscillations contribute to subjective time perception (Duration_TestReported_) across different conditions, we modelled single-trial reported duration using a GLMM with a Gamma distribution simultaneously including theta power, alpha power, their interaction with condition, and the aperiodic exponent (Supplementary Table 9). Because Compress was the only condition in which the instructed pace fell below the encoded duration, we predicted that the alpha–duration relationship would reverse sign under compressed replay. A planned orthogonal contrast on the alpha × condition interaction (Compress vs. the mean of Free and Expand) showed that the alpha–duration slope diverged under Compress. Specifically, the alpha–duration relationship during Compress trials differed from the average of the other conditions (*b* = 0.763, *z* = 3.109, *p* = .002) (Fig. 3ci). Simple slope analyses indicated that alpha power showed a positive but non-significant effect in predicting reported duration within the Compress condition (*b* = 0.357, *z* = 1.837, *p* = .066), while negative but non-significant trends were observed in the Free (*b* = -0.445, *z* = -1.584, *p* = .113) and Expansion (*b* = -0.367, *z* = -1.431, *p* = .152) conditions. The corresponding analysis of Duration_TestReplay_ is reported in Supplementary Figure 6b.

Conversely, theta activity appeared to show an opposing pattern, more closely aligning with the behavioural pattern, in which expanded replay lengthened reported duration. Because expanded replay should allow events to be re-segmented at a finer grain, we predicted a more positive theta–duration relationship under Expand. Within the same model, a planned orthogonal contrasts on the theta × condition interaction (Expand vs. the mean of Free and Compress) indicated a more positive theta–duration slope under Expand (*b* = 0.504, *z* = 2.693, *p* = .007) (Fig. 3cii). Follow-up simple slopes suggested a significant positive relationship between theta power and reported duration in the Expand condition (*b* = 0.446, *z* = 2.504, *p* = .012), whereas this relationship was non-significant but negative in the Free (*b* = -0.057, *z* = -0.361, *p* = .718) and Compress (*b* = -0.058, *z =* -0.451, *p* = .652) conditions.

Finally, neither alpha power nor theta power significantly predicted memory content, measured by gist accuracy (alpha: *b* = 0.253, *z* = 0.189, *p* = .850; theta: *b* = 0.492, *z* = 0.584, *p* = .559) or detailed recall (alpha: *b* = -0.144, *z* = -0.418, *p* = .676; *b* = 0.010, *z* = 0.053, *p* = .958). Thus, the alpha/theta power selectively tracks temporal processing rather than general memory content. Taken together, these results suggest that alpha oscillations may preferentially support the processing of compressed temporal experiences, while theta oscillations may be more engaged during temporal dilation.

### Replay speed-altered duration judgments cannot be explained by neural representations of memory content

Having shown that the replay speed selectively scales the retrospective reported duration without altering memory content at behavioural level, we further examined its effect at the neural representational level using RSA. Event-specific neural pattern similarity provides a fine-grained measure of content integrity, capable of detecting subtle representational shifts even when overt behavioral performance (e.g., recall or recognition) shows no apparent changes (Liu et al., 2021; Ritchey et al., 2013). Specifically, we calculated the diagonal mean of Manipulation–Test Similarity (MTS) to index the neural pattern similarity of memory representations between the instructed replay and the spontaneous replay at test (Fig. 4a). For each subevent, MTS was computed as within-subevent similarity (the diagonal) minus between-subevent similarity (the off-diagonal; see Methods).

**Fig. 4:**
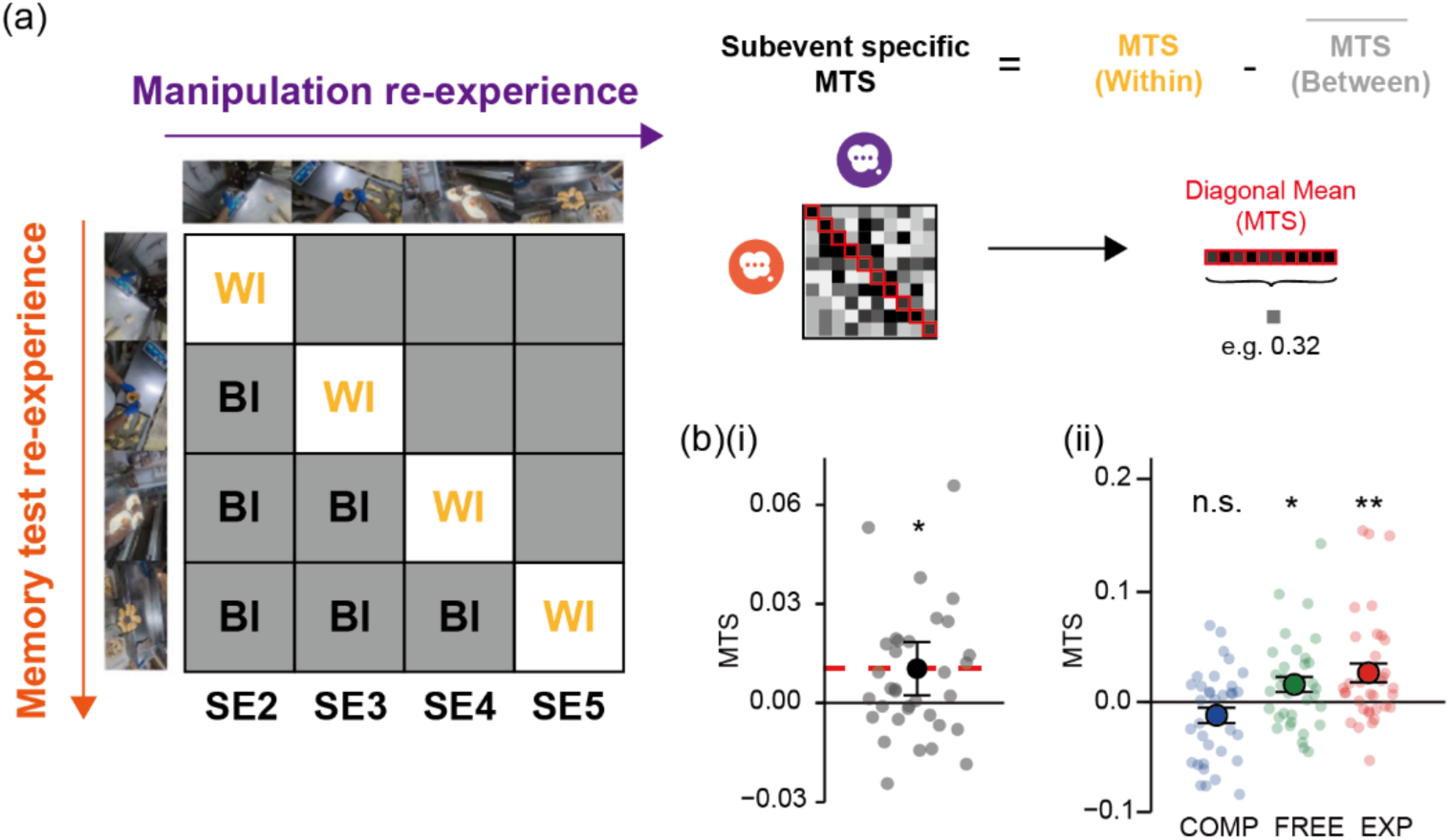
Oscillatory timing operates independently of memory content. (**a)** Schematic of Representational Similarity Analysis (RSA). The Manipulation-Test Similarity (MTS) index quantifies the neural reinstatement of subevent-specific information. It is calculated by subtracting the average similarity of distinct subevents (Between-Item; BI) from the similarity of the same subevent across Manipulation and Test phases (Within-Item; WI). (b) Neural reinstatement strength. **(i)** Aggregate MTS values relative to chance (zero line). Grey dots represent individual participant values. The red dashed line indicates the group mean. **(ii)** MTS values plotted by condition (Compress: blue, Free: green, Expand: red). Data are presented as mean values SEM. Transparent dots represent individual participant means.

Across conditions, MTS was reliably above zero (*t*(34) = 2.632, *p* = .013), confirming event-specific reinstatement at test (Fig. 4bi). By condition, reinstatement was significant for Expansion (*M* = 0.025, *SD* = 0.051, *t*(34) = 2.928, *p* = .006, *d* = 0.495) and Free replay (*M* = 0.015, *SD* = 0.042, *t*(34) = 2.204, *p* = .034, *d* = 0.372), but not for Compression (*M* = −0.010, *SD* = 0.043, *t*(34) = −1.373, *p* = .179). Reinstatement differed across conditions (repeated-measures ANOVA, *F*(2,68) = 5.185, *p* = .008; BF₁₀ = 12.60), being weaker after Compression than after Expansion (*b* = −0.035, *t*(68) = −3.120, *p* = .008; Holm-corrected), with a non-significant trend relative to Free replay (*b* = −0.026, *t*(68) = −2.251, *p* = .055) and no difference between Free and Expansion (*p* = .388). Thus, compressed replay was followed by weaker manipulation–test reinstatement, despite intact behavioural recall. Because MTS compares two phases, this reduction may partly reflect how the event was represented during compressed replay rather than at test alone.

Critically, we tested whether fluctuations in representational strength drove the observed shifts in retrospectively reported duration. According to the storage-size account (Ornstein, 1969), if these duration judgements rely on the volume of retrieved content, variations in MTS across subevents should predict reported duration. Importantly, adding trial-wise MTS into our existing Interaction Model of Duration_TestReported_ did not significantly improve model fit (*χ²*(1) = 3.106, *p* = .078), and MTS was not a significant predictor within the model (*b* = 0.013, *z* = 1.764, *p* = .078) (Supplementary Table 10).

Crucially, the condition-specific oscillatory effects on reported duration persisted when MTS was included as a covariate. The alpha × condition interaction remained significant (*χ²*(2) = 9.637, *p* = .008), as did both planned contrasts: the alpha–duration slope remained more positive under Compress than under the other conditions (*b* = 0.761, *z* = 3.104, *p* = .002), while theta showed the corresponding effect during Expand (*b* = 0.511, *z* = 2.733, *p* = .006). Bayesian model comparison yielded moderate-to-strong evidence that adding MTS did not improve the reported-duration model (*BF_01_* = 8.96). Causal mediation analysis further indicated that memory content did not mediate the oscillatory influence on reported duration, with a non-significant average causal mediation effect (ACME) for both alpha in the Compress condition (ACME = 0.925, 95% CI [−15.549, 18.222], *p* = .890) and theta in the Expand condition (ACME = −4.164, 95% CI [−34.232, 19.355], *p* = .734). Together, these results indicate that the oscillatory tracking of reported duration cannot be explained by the amount of event-specific content reinstated during replay.

## Discussion

The present study provides novel evidence that people can actively reconstruct memories about the temporal structure of naturalistic events. We demonstrated that manipulating mental replay speed carries over into subsequent retrospective duration judgments, while leaving recalled content unaffected. At the neural level, the relationship between oscillatory power during replay and later reported duration depended on the speed instruction. Under compressed replay, trials with higher alpha power tended to be judged as longer, whereas under free and expanded replay this relationship was absent or reversed. Conversely, under expanded replay, trials with higher theta power tended to be judged as longer, a relationship not observed under free or compressed replay. Critically, neither relationship was explained by the reinstatement of event-specific content. Although reinstatement itself varied with replay speed, it did not predict reported duration, did not improve model fit, and did not mediate the oscillatory effects. Remembered duration is therefore tracked by oscillatory dynamics that are dissociable from what is recalled, at both the behavioural and the neural representational level, which may allow the brain to adjust the evaluation of remembered time without altering the memory content.

### Dissociable temporal context and content of episodic memory

One of our main contributions is that the temporal judgment of a memory can be dissociated from its content. Although neural reinstatement of event was robust and varied with replay speed, it did not predict reported duration, challenging the storage-size hypothesis, in which remembered duration is read out from the amount of retained information (Ornstein, 1969). Content and time were thus not fully independent, as reinstatement itself varied with replay speed; rather, they dissociated specifically at the level of explicit duration judgments. The temporal readout of a memory can therefore be re-paced, compressed for rapid review or expanded for closer inspection, while its underlying representation continues to carry event-specific content (Lositsky et al., 2016).

We suggest this dissociation reflects a functional division between two properties of a retrieved memory that can be modified independently, a division that bears on competing accounts of how episodic memory is organized. Integrative models hold that an event and its temporal context form a single bound representation, such that retrieval recovers a drifting context signal that is itself part of the trace and that remembered duration is inferred from the volume of content recalled (Ornstein, 1969). Such accounts predict that temporal pacing cannot be adjusted without disturbing content, which our findings do not support. Independent-storage accounts, by contrast, posit a temporal code stored separately from sensory and semantic content, as suggested by time cells and ramping cells that track elapsed time irrespective of what is experienced (Reddy et al., 2021; Tsao et al., 2018; Umbach et al., 2020); yet reinstatement in our data varied with the speed instruction, which a fully separate code would not predict.

These findings inform two contrasting accounts of how remembered duration is derived. Under the storage-size account, and the related integrative models in which an event and its temporal context are encoded as a single bound trace (Howard & Kahana, 2002; Polyn et al., 2009), remembered duration is inferred from the quantity of retrieved content (Ornstein, 1969) or from the contextual change that content spans (Lositsky et al., 2016). This view predicts that the remembered duration of an event cannot be altered without a corresponding change in how much of it is retrieved. Our results contradict this prediction. Specifically, replay speed shifted reported duration without affecting recall, and trial-wise reinstatement neither predicted reported duration nor mediated the oscillatory effects. By contrast, the independent temporal storage account, which proposes that temporal structure is represented separately from sensory and semantic content (Reddy et al., 2021; Tsao et al., 2018; Umbach et al., 2020), readily accommodates this dissociation. This account, however, cannot explain the weaker reinstatement observed after compressed replay, because fully separate storage implies that re-pacing an event should leave reinstatement of its content unaffected. Taken together, our findings support independent temporal storage at the level of duration judgments, while arguing against a strict separation of time and content at the level of neural representation.

One account that reconciles these observations is that time and content are linked within the memory representation but can be accessed and modified separately at retrieval. Retrieval renders a memory temporarily labile and open to modification (Hupbach et al., 2008; Nadel et al., 2000), and distinct features of an event, such as specific items and their contexts, can be recalled, forgotten and updated independently (Brady et al., 2013; Yonelinas, 2002). Remembered duration may constitute one such attribute, such that an episode’s temporal pacing can be recalibrated while its content is preserved. This flexibility is mechanistically plausible, as hippocampal and entorhinal circuits represent temporal structure as a tunable sequence that is partially dissociable from event content (Buzsáki & Tingley, 2018; Tsao et al., 2018; Whittington et al., 2020). Such independent control over the temporal metric may also support learning and future planning; adaptive prospective simulation requires the cognitive flexibility to scan, compress and generalize past experiences rather than strictly re-experiencing them in veridical time (Richards & Frankland, 2017; Schacter et al., 2007). The framework further predicts the converse dissociation, in which episodic content is altered while remembered duration is preserved, a hypothesis that remains to be tested empirically.

### The Oscillatory Gating of Subjective Time

Alpha power was associated with reported duration, consistent with a role in gating whether the encoded timeline is updated during replay. Higher alpha power reflects a closed gate that inhibits updating and shields the maintained representation, whereas lower power reflects an open gate that permits it (Jensen & Mazaheri, 2010; Klimesch, 2012; Manza et al., 2014). The condition-level pattern fits this account: alpha power was lowest in Compression, where updating the encoded pacing was most successful, and higher where replay left the representation more intact. Within each instruction, higher trial-wise alpha predicted shorter reported durations under Free and Expanded replay but longer durations under Compression. This gating role for alpha is not specific to temporal memory; the same principle governs the updating of episodic content more broadly. When a consolidated memory is reactivated, it becomes labile and open to revision, and the degree of updating tracks alpha power: lower alpha during and after reactivation predicts greater integration of new information into the existing memory (Zhu et al., 2019). Alpha desynchronization likewise accompanies the reinstatement of stored representations into awareness during retrieval, consistent with reduced alpha releasing memory content for processing (Griffiths et al., 2019; Hanslmayr et al., 2012). Notably, alpha during manipulation did not predict MTS under Compression (*p* = .993), suggesting that any such gating operated on the temporal metric rather than on the reinstatement of event features, which MTS indexes. Future studies using simultaneous EEG–fMRI, intracranial recordings or source-resolved MEG could localize where this gating occurs and test whether it acts selectively on temporal rather than featural content. Within working memory, alpha similarly gates which representations are updated versus protected from interference (Manza et al., 2014). Across these domains, lower alpha marks a state in which the stored representation is open to revision, and higher alpha a state in which it is shielded — the same disinhibition-permits-updating logic we observe for the remembered duration, here applied to its temporal rather than its featural content.

### Theta Oscillations and the Expansion of Time

Theta oscillations (4–8 Hz) offer a candidate mechanism for the lengthening of subjective time under expanded replay. Theta activity has long been implicated in the temporal organization of memory, supporting the encoding of event order and the binding of successive elements into a coherent episodic sequence (Heusser et al., 2016; Hsieh et al., 2014; Lisman & Jensen, 2013). Consistent with this role, the relationship between theta power and reported duration was more positive under expanded than under free and compressed replay, such that trials with higher theta power tended to be judged as longer. One interpretation is that theta activity acts as an internal pacer, marking successive points in the unfolding of an episode. During expanded replay, the same episodic content is traversed more slowly, allowing additional theta cycles to accumulate within its span and thereby impose finer temporal subdivisions between encoded elements. Remembered duration would consequently lengthen not because more content is retrieved, but because the episode is partitioned into a greater number of internal intervals.

This account parallels spatial navigation, in which hippocampal theta supports the integration of a traversed path and scales with the distance covered (Bellmund et al., 2018; Buzsáki & Moser, 2013; Vass et al., 2016). Expanded replay may similarly accumulate more theta cycles to span a greater temporal distance, an extent that is temporal rather than informational, such that a longer trajectory need not carry more retrievable detail. This interpretation is consistent with our observations that theta power predicted neither gist accuracy nor detail recall, and that its relationship with reported duration was not explained by reinstatement. Theta may therefore index the temporal extent of replay rather than its content, although this interpretation remains tentative. Future studies that directly manipulate segmentation grain could test whether theta activity scales with the number of event boundaries independently of replay speed.

### Limitations and Future Directions

Several limitations of the present work suggest avenues for future research. First, we did not collect a duration judgment before the replay manipulation, since asking for one would have disclosed the timing task and invited prospective strategies such as counting (Block & Zakay, 1997). Second, while theta oscillations tracked temporal expansion, scalp EEG cannot definitively localize these signals to the hippocampus; future studies utilizing MEG or intracranial recordings are required to map the specific hippocampal-neocortical dynamics supporting this scaling. Finally, these findings offer a novel framework for understanding clinical disorders characterized by temporal distortion. For instance, the distorted time perception in Alzheimer’s disease (El Haj et al., 2013; El Haj & Kapogiannis, 2016b) and the overestimation of durations reported in post-traumatic stress disorder (Vicario & Felmingham, 2018) may stem from specific disruptions in these oscillatory timing mechanisms, dissociable from the broader memory deficits often assumed to drive these symptoms.

## Conclusion

In summary, the duration of an event is not recorded passively but reconstructed at retrieval, and this reconstruction is accompanied by distinct oscillatory signatures. The relationship between alpha power and reported duration depended on the instructed replay speed, diverging in direction between compressed replay and free or expanded replay, consistent with a role in gating whether the encoded timeline is updated. Theta showed a complementary, exploratory link to temporal expansion. Critically, neither effect was explained by the content of the retrieved memory, indicating that the subjective duration of an experience can be rescaled without altering what is remembered. This capacity to re-pace a memory while preserving its content may serve the flexible use of past experience in learning and future planning.

## Methods

### Participants

Fifty-seven individuals (45 female; Mage = 24.1 years, SD = 2.59 years) were recruited from the University of Hong Kong. All participants had normal or corrected-to-normal vision and hearing without color blindness, and none had chronic medical conditions, severe mental health history, neurological disorders, or active psychiatric diagnoses. Sixteen participants were excluded: two for falling asleep during the task and fourteen for failing the manipulation check (i.e., mean manipulation replay durations not conforming to the expected Compress < Free < Expand pattern), yielding a final sample of 41. An a priori power analysis (G*Power 3.1.9.7; α = 0.05, power = 0.80, r = 0.50) indicated a minimum of 28 participants was required to detect a conservative a priori effect size of η² = 0.06. This estimate was chosen to account for effect size inflation (Szucs & Ioannidis, 2017), despite larger effects reported in related temporal memory research (Bangert et al., 2019; Jeunehomme & D’Argembeau, 2019; Morris & Fritz, 2013). Participants provided written informed consent and received HKD 400 (∼USD 51) compensation. This research was approved by the University of Hong Kong Human Research Ethics Committee for Non-Clinical Faculties (EA210341).

### Materials

Fifteen videos (175 seconds each) were created by selecting and combining first-person perspective segments from the Ego4D dataset, which closely approximate natural event experience (Grauman et al., 2022) (See the list of videos used in Supplementary Table 1). Each video depicted a hierarchical overarching event comprising five distinct subevents with clear boundaries (e.g., kneading dough, washing eggs in the ‘Bread’ video; Fig. 1B). To ensure clear boundaries, subevents were selected based on two criteria: (1) activities had to be readily distinguishable within the time constraints; and (2) segments had to include recognizable actions and objects describable without technical expertise. The first subevent (15s) served as a reference segment for speed manipulations. To prevent predictable temporal patterns, the remaining four subevents (two 30s, two 50s) were presented in a randomized order. The 15 videos were randomly and equally distributed across three experimental conditions (5 videos each), with two additional videos used for practice.

Each subevent was filmed in a single continuous shot with no manual editing, visual effects, or subtitles to maintain natural continuity. Subevents were trimmed and combined into complete sequences, then exported at 1920 × 1080 resolution. To validate the stimuli, three independent coders completed a boundary identification task; we retained only the videos where all three coders successfully identified the subevent transitions within 5 seconds of the actual boundaries. The experiment was conducted using PsychoPy version 2023.1.3. Verbal recall during retrieval was recorded using a Sony Stereo IC Recorder (ICD-UC570F).

### Procedure

The study employed a within-subjects design with three mental replay conditions (Compress, Free, Expand) across three phases: encoding, manipulation, and testing. Pre-experiment questionnaires, including demographic information, the Pittsburgh Sleep Quality Index (PSQI; Buysse et al., 1989), the Insomnia Severity Index (ISI; Bastien & Morin, 2001), and the Depression Anxiety Stress Scales (DASS; Lovibond & Lovibond, 1995). Participants reported low scores on all measures, within the normal range for depression (*M* = 5.024, *SD* = 6.421), anxiety (*M* = 5.122, *SD* = 4.496) and stress (*M* = 7.463, *SD* = 6.797) on the DASS-21, and below the threshold for clinically significant insomnia on the ISI (*M* = 6.488, *SD* = 4.996). These measures were therefore not considered in further analyses.

#### Encoding phase

During encoding, participants viewed each video as if personally experiencing the event. Each video was shown only once to mirror real-world conditions, where events are typically experienced without repetition. While viewing, participants pressed the spacebar whenever they noticed a transition between subevents. This active segmentation served two purposes: (1) it confirmed that participants recognized subevent transitions, allowing us to determine the actual duration of each recognized subevent, and (2) it helped maintain attention throughout the video.

Participants expected a memory test but were kept naïve to the duration estimation task to prevent covert timing strategies, such as counting. Videos were presented in a randomized order, each preceded by a targeted retrieval word cue (e.g., “Bread”). After each video, participants indicated whether they had previously seen the content. They then completed a brief arithmetic distractor task (three problems; M = 9.95s, SD = 3.50s) to prevent immediate, spontaneous mental replay.

#### Manipulation phase

Immediately following the distractor task, participants mentally replayed the entire event sequentially and without skipping content. Videos were counterbalanced across three conditions: Expand (0.5× speed), Compress (4× speed), and Free (self-paced control). The 4× rate was chosen because mental replay naturally occurs faster than real time (Denovellis et al., 2021; Jeunehomme et al., 2020; Michelmann et al., 2019).

Trials began with a 3-second countdown and cue (“Video” for Expand/Compress; “Replay” for Free). To serve as a temporal guide, the first subevent was re-presented at the target speed in the Expand and Compress conditions. Participants then mentally replayed subevents 2–5 at this anchored pace. Crucially, the Free condition omitted this calibration segment to prevent anchoring participants’ internal clocks and contaminating their natural, unconstrained replay tendencies.

Across all conditions, participants pressed the spacebar after mentally replaying each subevent to verify timing adherence. To maintain experimental consistency, the first subevent was excluded from all subsequent analyses. Finally, participants rated their replay quality and completed a temporal order task—matching five screenshots to their respective subevents—to verify structural retention (Fig. 1c).

#### Memory test phase

All 15 videos were tested in randomized order, with duration and memory evaluated subevent by subevent.

For each subevent, participants first indicated recall success (Yes/No). To determine the effect of the speed manipulation on retrospective duration judgment, participants then estimated the subevent’s original duration in seconds via direct numerical entry (15-second response window).

Next, participants mentally replayed the subevent as vividly as possible at their natural pace, with explicit instruction that they need not follow the manipulation speed but use their normal speed. Replay duration was recorded via spacebar press. To assess whether the manipulation affected memory content in addition to temporal aspects, participants then provided detailed verbal recall for up to one minute, describing all remembered details including characters, items, and contextual elements.

Finally, participants rated the percentage of content remembered and replay vividness to capture subjective memory quality, and repeated the temporal order task to verify retention of event structure.

#### Total duration assessment

Following the completion of all subevent assessments, a separate test was administered for whole-event duration. Participants were presented with each video’s word cue and estimated the total duration of that complete video. This measure examined whether subevent-level effects extended to whole-event duration judgments.

### Verbal recall analysis

#### Preprocessing

Audio recordings were segmented by subevent (Audacity) and transcribed using OpenAI’s Whisper model (medium variant). Transcriptions underwent manual review, supplemented by ChatGPT-4 to identify and correct homophones and semantic inconsistencies common in automated speech recognition outputs.

#### Gist Accuracy Coding

To establish a reference standard for gist accuracy, we asked human coders to view each video and identify the core gist of each subevent, defined as the essential action and its result. This produced scoring templates against which participants’ recall could be evaluated.

To assess gist accuracy, we used ChatGPT-4 to summarize each participant’s recall of each subevent and to rate its semantic similarity to the corresponding template. This approach focused on meaning rather than exact word matching, accepting variations including synonyms (e.g., “pasta” vs. “noodles”), paraphrasing, and action-outcome equivalence (e.g., “nailing wood together” vs. “using nails to fix wooden boards”), while excluding superficial attributes such as colors, quantities, and object positioning. Similarity scores ranged from 0 (no relevant match) to 1 (complete capture of key meaning). Scores were then converted to binary gist scores using a 0.5 threshold.

#### Detail Content Coding

To assess recall richness, human coders comprehensively cataloged all identifiable video elements into five predefined categories: character, action, environment, central object, and peripheral object. The number of correctly recalled details was quantified by tallying matches between participants’ transcriptions and this master template with ChatGPT-4. Matching criteria accepted synonymous phrases, minor misspellings, and general terms for specific labels (e.g., “meat” for “chicken breast”), focusing on core identity rather than superficial attributes.

#### Reliability Validation

To validate the automated NLP scoring approach, a human rater independently scored a subset of the data (the first five participants) using identical criteria. High agreement was achieved for both gist accuracy (94.4%) and detail content (96.6%), validating the use of ChatGPT-4 evaluations for all final statistical analyses.

### Behavioral Data Analysis

All statistical analyses were performed using R (version 4.3.0; R Core Team, 2023) with packages including dplyr (Wickham et al., 2023), ggplot2 (Wickham, 2016), lme4 (Bates et al., 2015), lmerTest (Kuznetsova et al., 2017), emmeans (Lenth, 2023), car (Fox & Weisberg, 2019), and mediation (Tingley et al., 2014). At the participant level, those whose manipulation duration did not follow the expected order (COMP < FREE < EXP) were excluded. Following the initial behavioral manipulation exclusions, univariate outliers exceeding 1.5 times the interquartile range (IQR) were removed, resulting in the exclusion of two participants from test replay duration analyses and two from retrospective duration analyses. At the trial level, 42 individual trials were excluded because participants failed to identify all five subevents during initial encoding.

Temporal precision during replay was quantified as the percentage of spacebar presses occurring within a 5-second window of true subevent boundaries, explicitly penalizing indiscriminate pressing. Differences across conditions were evaluated using a non-parametric Friedman test followed by Bonferroni-corrected Wilcoxon signed-rank tests. To accommodate the positively skewed distribution of continuous timing variables (spontaneous replay and retrospective duration), we employed Generalized Linear Mixed Models (GLMMs) utilizing a Gamma distribution and log link function. Random intercepts were included for both participants and specific events. To disentangle the inherently collinear effects of the categorical manipulation and the actual manipulated duration experienced, we constructed two separate GLMMs for each behavioral outcome. The Condition Model evaluated the categorical fixed effect of the experimental group (Compress, Free, Expand) (Supplementary Table 2 and 6), whereas the Experience Model evaluated the continuous fixed effect of the participant’s specific, Z-scored manipulated duration (Supplementary Table 3 and 7). Both models controlled for the actual original duration of the subevent (30s vs. 50s).

### EEG acquisition and preprocessing

Continuous EEG was recorded at 500 Hz using an eego amplifier (ANT Neuro, The Netherlands) and a 61-channel gel-based EEG cap (Waveguard™ original) configured to the International 10–20 system (CPz reference, AFz ground). Impedances were maintained below 20 kΩ. Additional electrodes were placed on the bilateral mastoids (M1, M2) and 1.5 cm left of the external canthus for EOG monitoring.

EEG data were processed offline using MATLAB (MathWorks Inc., version R2024A) and EEGLAB (version 2024.1)(Delorme & Makeig, 2004). Auxiliary channels (EOG, M1, and M2) were removed, and data were downsampled to 250 Hz. The data were bandpass-filtered (0.1–40 Hz) and notch-filtered at 50 Hz to remove line noise. Channel locations were assigned using the standard 10–5 electrode montage. Bad channels were identified through visual inspection, removed, and interpolated using spherical interpolation to maintain the original channel configuration.

Prior to Independent Component Analysis (ICA), data were segmented by task phase (encoding epochs matched video duration; replay epochs utilized the longest trial) with a ∼10s buffer appended to prevent edge effects. Following extended Infomax ICA (Pion-Tonachini et al., 2019), components classified via ICLabel (>90% probability of ocular or muscular artifacts) were removed. Cleaned data were re-referenced to the grand average, and epochs exceeding ±200 μV were automatically rejected.

Six participants were excluded from EEG analyses due to insufficient data quality (<5 usable epochs per condition) but retained for behavioral analyses, yielding a final EEG sample of 35 participants.

### Time-frequency resolved EEG Power Spectral Analysis

Manipulation and test replay epochs were first downsampled to 100 Hz prior to Morlet wavelet decomposition (FieldTrip; 1–40 Hz in 1 Hz steps, 5-cycle width, Gaussian width of 3). Epochs shorter than 2 seconds were excluded; given the original 30–50s subevent durations, such brief mental replays reflect extreme compression rates that far exceed the maximum instructed limit (4x) and are therefore incompatible with our manipulation instructions. This threshold removed five total trials (Free: three trials; Compress: two trials).

Power spectral densities were estimated using Welch’s method and parameterized using the Fitting Oscillations & One Over F (FOOOF) package (Donoghue et al., 2020) implemented in Jupyter Notebook. Because replay duration varied across subevents and participants, a sliding window approach was adopted to ensure consistent temporal sampling. Spectral densities were computed using 2-second windows with 50% overlap, applying Welch’s method with Hamming windowing across the 1–40 Hz frequency range. These spectral densities were temporally averaged across sliding windows within each subevent trial, and then spatially averaged across EEG channels. Finally, FOOOF models were fit to each participant’s subevent trials across all conditions, with the knee parameter fixed at k = 1.0 to ensure identical fitting procedures and prevent artificial differences in oscillatory power estimates.

### Neural Oscillatory Modeling and Model Selection

To evaluate whether neural oscillations during the manipulation phase predicted subsequent temporal memory, we constructed Generalized Linear Mixed Models (GLMMs; Gamma family, log link). Base models predicted temporal memory from spectral features (theta power, alpha power, and the aperiodic exponent), controlling for original video duration and experimental condition, with random intercepts for Subject and Event. Model selection via Likelihood Ratio Tests (LRT) compared these base models against two alternatives: a Full Spectrum Model incorporating delta (1–3 Hz), beta (13–30 Hz), and gamma (30–40 Hz) bands, and an Interaction Model including frequency-by-condition interaction terms (Supplementary Table 5 and 9). The models underlying the main results were specified as follows, with Subject and Event entered as random intercepts throughout:

**Band Power Model**: *band ∼ Condition + (1 | Subject)*

**Base Model**: *ReportedDuration ∼ VideoLength + Condition + theta + alpha + aperiodic exponent + (1 | Subject) + (1 | Event)*, Gamma(log).

**Interaction Model**: *ReportedDuration ∼ VideoLength + alpha × Condition + theta × Condition + aperiodic exponent + (1 | Subject) + (1 | Event)*, Gamma(log).

**Full Spectrum Model**: the Interaction Model with delta, beta, and gamma power added.

Planned contrasts (Compress versus the mean of Free and Expand on the alpha × condition term; Expand versus the mean of Free and Compress on the theta × condition term) were computed on the fitted interaction slopes using estimated marginal trends.

We included theta and alpha power, and the aperiodic exponent simultaneously in these models to rigorously isolate their unique predictive contributions. Additionally, including the aperiodic exponents ensures that targeted theta/alpha oscillations are not artifacts of global broadband shifts. To ensure our findings were not driven by model complexity or collinearity, we conducted supplementary sensitivity analyses. We modelled theta and alpha power individually for each condition. These separate, independent models yielded identical patterns of significance with numerical differences, confirming the robustness of our integrated modelling approach.

Simultaneous inclusion of theta, alpha, and the aperiodic exponent rigorously isolated their unique predictive contributions while confirming that targeted oscillations were not mere artifacts of global broadband shifts. Supplementary sensitivity analyses, modelling theta and alpha individually per condition, yielded identical significance patterns, confirming the robustness of this integrated modelling approach. Finally, all predictors exhibited acceptable multicollinearity across all models (Generalized Variance Inflation Factors < 2.25, equivalent to VIF < 5; Supplementary Table 2-4, 6-8).

### Episodic Content Memory Modelling

To evaluate whether neural oscillatory states predicted subsequent episodic recall, gist accuracy (binary) and the total count of recalled details were assessed as a function of theta power, alpha power, and the aperiodic exponent. These predictive relationships were quantified using GLMM, specifically, a binomial model (logit link) for gist accuracy and a Poisson model (log link) for detailed recall. Both models evaluated the spectral features simultaneously to isolate their unique contributions, controlling for actual video length and experimental condition, with random intercepts included for Subject and Event. The aperiodic exponent was included as a covariate because it captures broadband (1/f) changes in the power spectrum, which are thought to reflect shifts in excitation–inhibition balance (Donoghue et al., 2020; Gao et al., 2017); controlling for it ensures that any association between band power and recall reflects oscillatory activity rather than broadband spectral change. The models were specified as follows.

**Gist accuracy**: *GistAccuracy ∼ alpha + theta + Condition + aperiodic exponent + VideoLength + (1 | Subject) + (1 | Event)*, binomial(logit).

**Detail recall**: *DetailCount ∼ alpha + theta + Condition + aperiodic exponent + VideoLength + (1 | Subject) + (1 | Event)*, Poisson(log).

### Representational similarity analysis

To evaluate the reinstatement of memory representations, we conducted representational similarity analysis (RSA) on EEG spectral power, comparing the manipulation and test replay phases. Spatial-spectral feature vectors were extracted using a sliding window (1.0s, 80% overlap) by temporally averaging power (1–40 Hz) and vectorizing across channels and frequencies. Cross-temporal similarity was computed using Spearman’s rank correlation, yielding a Fisher *Z*-transformed correlation matrix per participant and condition.

We calculated Manipulation-Test Replay Similarity (MTS) for each subevent by subtracting between-subevent similarity from within-subevent similarity. Specifically, for a target memory (e.g., Subevent 1), within-subevent similarity was defined as the correlation between its manipulation replay (M1) and corresponding test replay (T1) for the same subevent. Between-subevent similarity was computed by averaging correlations between the target subevent and all non-matching subevent across both phases (e.g., M1–T2 through M1–T60, and M2–T1 through M60–T1).

To evaluate statistical reliability, this index was subjected to two group-level tests addressing different questions: whether the effect generalizes across participants, and whether its magnitude exceeds what the data would produce under a null labelling. First, participant-level MTS was tested against zero with a one-sample two-tailed t-test (*t*(34) = 2.632, *p* = .013), treating participant as the unit of inference and verifying that within-subevent similarity exceeded between-subevent similarity across the cohort. Second, because the numbers of within- and between-subevent pairs entering MTS are unbalanced by design, we compared the observed value against a surrogate null distribution: on each of 1,000 permutations, subevent labels were shuffled across test-phase epochs while manipulation-phase epochs were held fixed, MTS was recomputed, and values were averaged across participants. The observed group mean was expressed as a z-score relative to this distribution, with p given by the proportion of surrogate means at or above it (*z* = 2.472, *p* = .005). This test was evaluated one-tailed because reinstatement predicts a strictly directional effect, with values below the null carrying no interpretation.

### Independence of Duration Judgments and Content Representations

#### GLMM

To investigate whether the associations between neural oscillatory activity and subjective duration judgments persisted after accounting for memory content reinstatement, we expanded our final GLMMs to include the MTS (*Z*-scored) as an additional fixed effect.

**Independence Model**: ReportedDuration ∼ VideoLength + theta × Condition + aperiodic exponent + alpha × Condition + MTS + (1 | Subject) + (1 | Event), Gamma(log).

We evaluated the unique predictive contribution of memory content via Bayesian Model Comparison, utilizing likelihood ratio tests and Bayes Factors (*BF_01_*) derived from the Bayesian Information Criterion (BIC) to compare the Independence Model (adding MTS) against the corresponding interaction model containing only oscillatory predictors (Supplementary Table 10).

To test the robustness of the condition-specific pattern, we analyzed estimated marginal trends (*emtrends*) from the Independence Models. Follow-up contrasts were computed to test whether the alpha slope in the Compress condition and the theta slope in the Expand condition remained statistically distinct from other conditions, even when explicitly controlling for MTS.

#### Mediation Analysis

To test whether MTS mediated the relationship between oscillatory power and reported duration, we conducted condition-specific mediation analyses using the mediation package in R (Tingley et al., 2014). Based on the profiles observed in the interaction models, we tested two specific pathways: (1) whether MTS mediated the effect of alpha power on Duration_TestReported_ in the Compress condition, and (2) whether MTS mediated the effect of theta power on Duration_TestReported_ in the Expand condition. To obtain stable estimates of representational similarity and to complement the trial-level analysis, mediation was assessed at the participant level: for each pathway, variables were averaged within participants for the relevant condition and entered into two linear regression models:

Mediator Model (Path A): Predicting MTS from the target oscillatory measure, controlling for manipulation duration.

Outcome Model (Path B): Predicting reported duration from both the target oscillatory measure and MTS, controlling for manipulation duration.

The indirect effects were estimated using Quasi-Bayesian Monte Carlo approximation with 1,000 simulations. Statistical significance was determined if the 95% confidence intervals (CI) for the Average Causal Mediation Effect (ACME) did not include zero.

## Acknowledgements

The authors gratefully acknowledge the valuable contributions of all participants involved in recruitment. The research was supported by The University of Hong Kong University Research Committee Internal Seed Fund for Basic Research to X. H.; Guangdong Basic and Applied Basic Research Foundation (No. 2024A1515012667 and No. 2023A1515110311) to J. L. This work was also supported by the grant from the Research Center for Brain Cognition and Human Development, Guangdong, China (No. 2024B0303390003).

## Data and code availability

The pre-processed data and custom code used to generate the main findings of this study are available in the OSF: https://osf.io/8ta3d/overview?view_only=dbfa161d95784cf890b379f7d6a6c1a6

## Author contributions

W. W. Y. Y., J. L., and X. H. conceived the study. W. W. Y. Y. collected the data. W. W. Y. Y., Z. M., Z. Y., D. C., Y. Z., J. L., and X. H. performed the analysis. W. W. Y. Y., J. L., and X. H. wrote the manuscript. W. W. Y. Y., Z. M., Z. Y., D. C., and Y. Z. revised the manuscript.

## Supplementary Figures and Tables

**Supplementary Figure 1.**
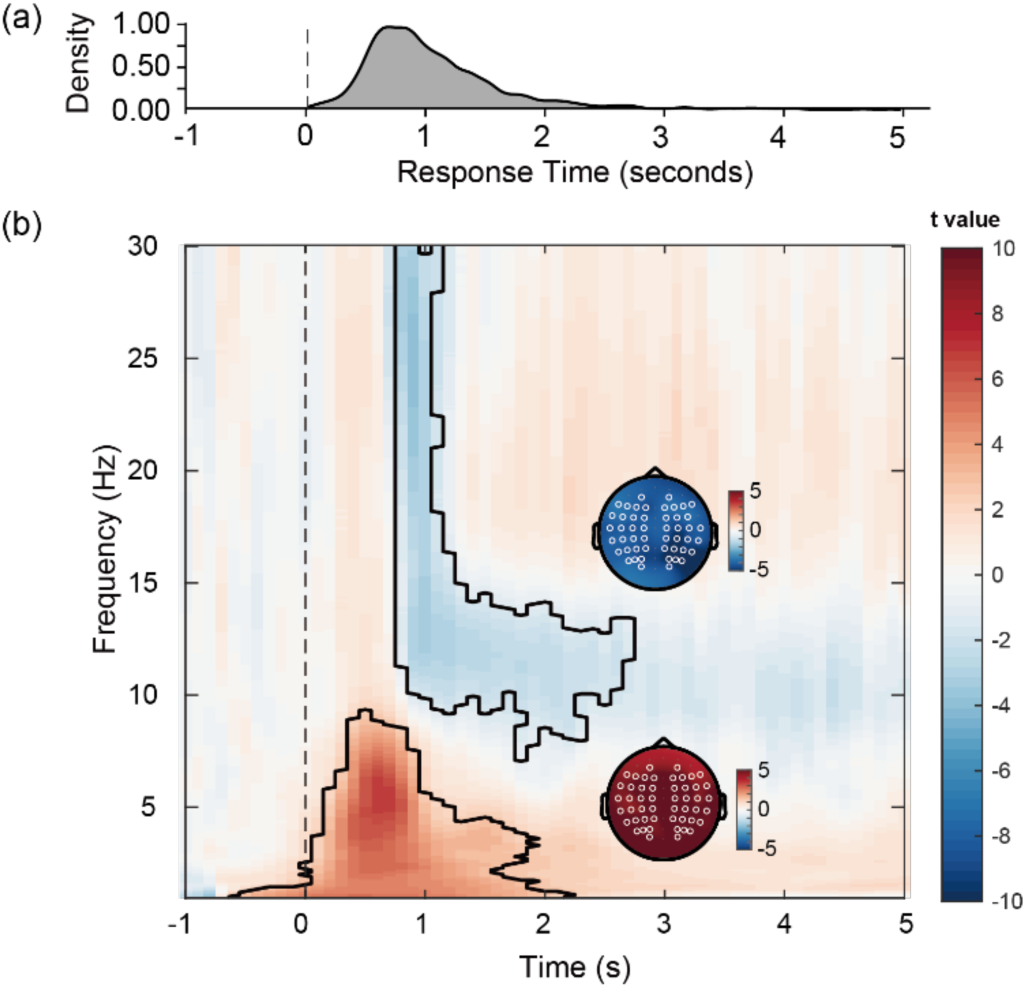
(a) Probability density function of response times for all boundary detection, with peak detection latency around 1 second post-boundary (dashed vertical line indicates boundary onset at 0 seconds). Peak response times were also consistent across Compress (*M* = 0.947 s), Expand (*M* = 0.904 s), and Free (*M* = 0.926 s), *F*(2, 80) = 1.004, *p* = .371. (b) Time-frequency analysis of EEG activity time-locked to event boundaries during video watching. The spectrogram shows statistical t-values (color scale) across frequencies (1-30 Hz). Dashed vertical line indicates boundary onset. The contour highlighted significant clusters.

**Supplementary Figure 2.**
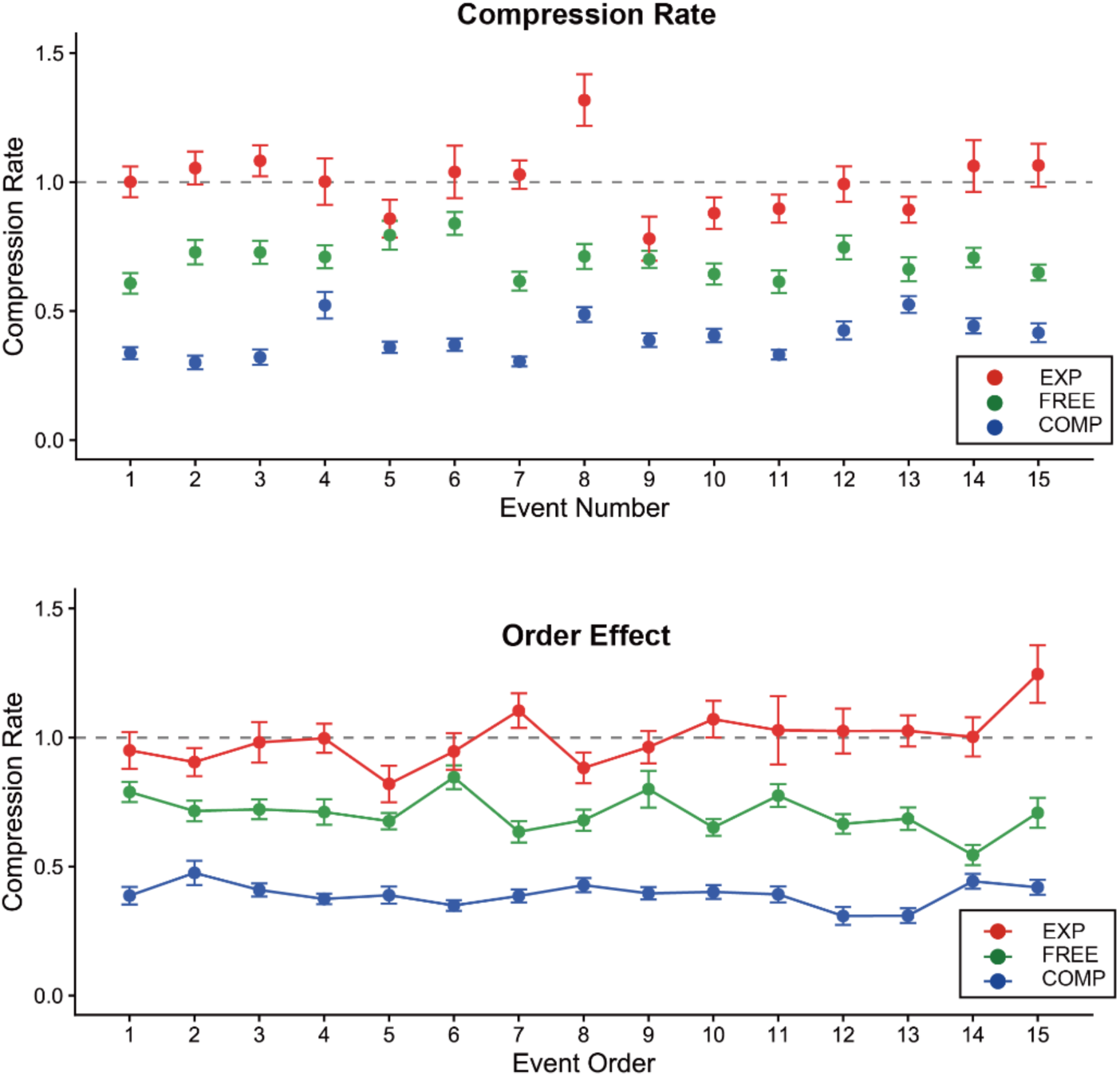
(a) Compression rate for each event during mental re-experience across the 15 events comprising each video. Compression rate is calculated as Duration_Mani_ / Duration_ActualVideo_ (175s). Values < 1 indicate replay shorter (faster) than the actual video; values > 1 indicate replay longer (slower) than the actual video. Colored dots show group means; error bars show ±1 SE. (b) Order effect showing manipulation duration for each order position during manipulation re-experience. Points show group means; error bars show ±1 SE. Compress: blue, Free: green, Expand: red.

**Supplementary Figure 3.**
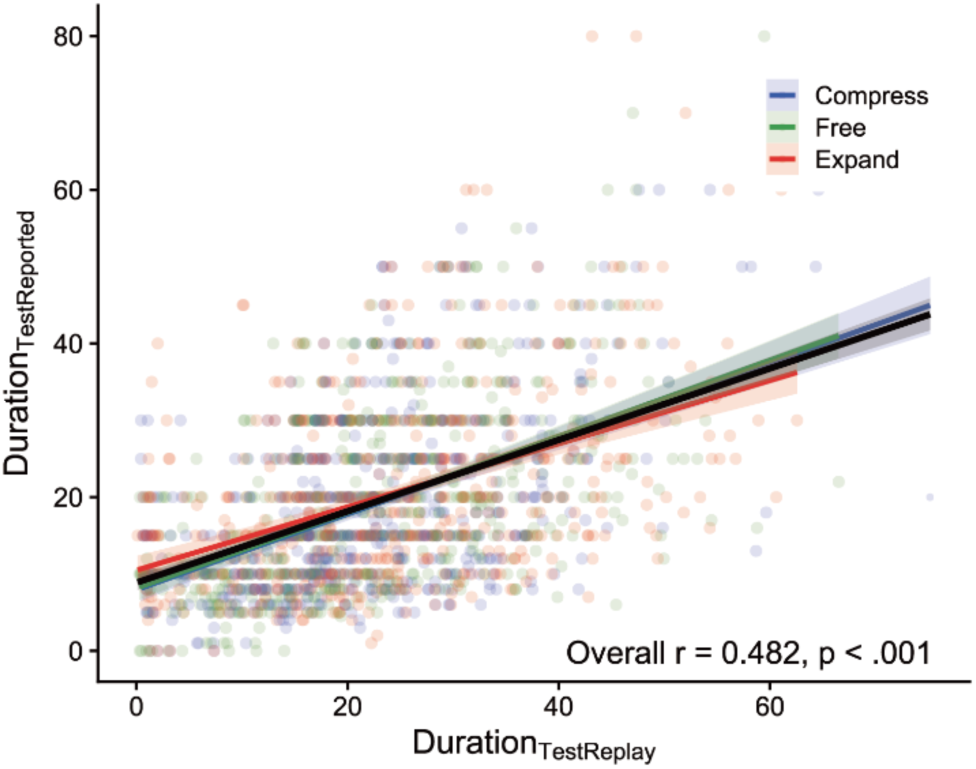
Correlation between Duration_TestReplay_ and Duration_TestReported_. The solid black line indicates the overall model fit; colored lines represent condition-specific linear fits with shaded areas denoting 95% confidence intervals. Transparent dots represent individual data points. Compress: blue, Free: green, Expand: red.

**Supplementary Figure 4.**
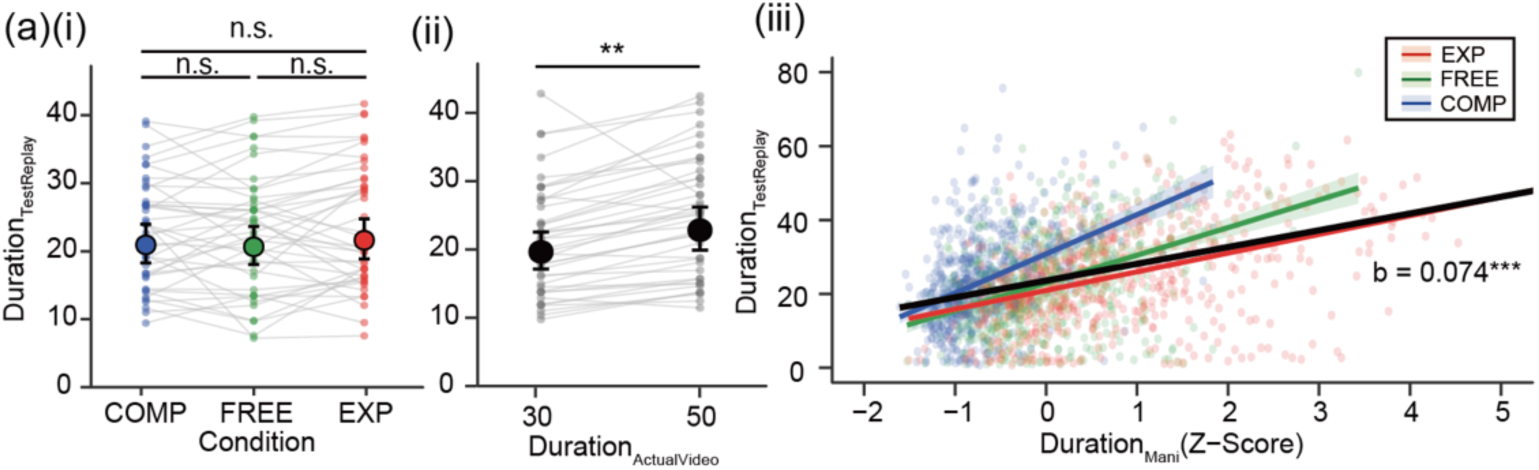
(a) Spontaneous replay duration during the test phase (Duration_TestReplay_). The speed manipulation did not significantly influence Duration_TestReplay_, *χ²*(2) = 2.02, *p* = .364 (i) Duration_TestReplay_ plotted across conditions. (ii) Duration_TestReplay_ by Duration_ActualVideo_ (30s vs. 50s). Solid dots represent model-estimated marginal means; error bars indicate 95% confidence intervals. (iii) Relationship between Duration_Mani_ (Z-scored) and subsequent Duration_TestReplay_. The continuous Duration_Mani_ predicted Duration_TestReplay_ (*b* = 0.074, *z* = 4.454, *p* < .001). The solid black line indicates the model fixed effect; colored lines represent condition-specific fits with shaded areas denoting 95% CI. Transparent dots represent individual participant means, connected by grey lines in (a) and (b). Compress: blue, Free: green, Expand: red. \**p* < .05, \*\*\**p* < .001, n.s. = not significant.

**Supplementary Figure 5.**
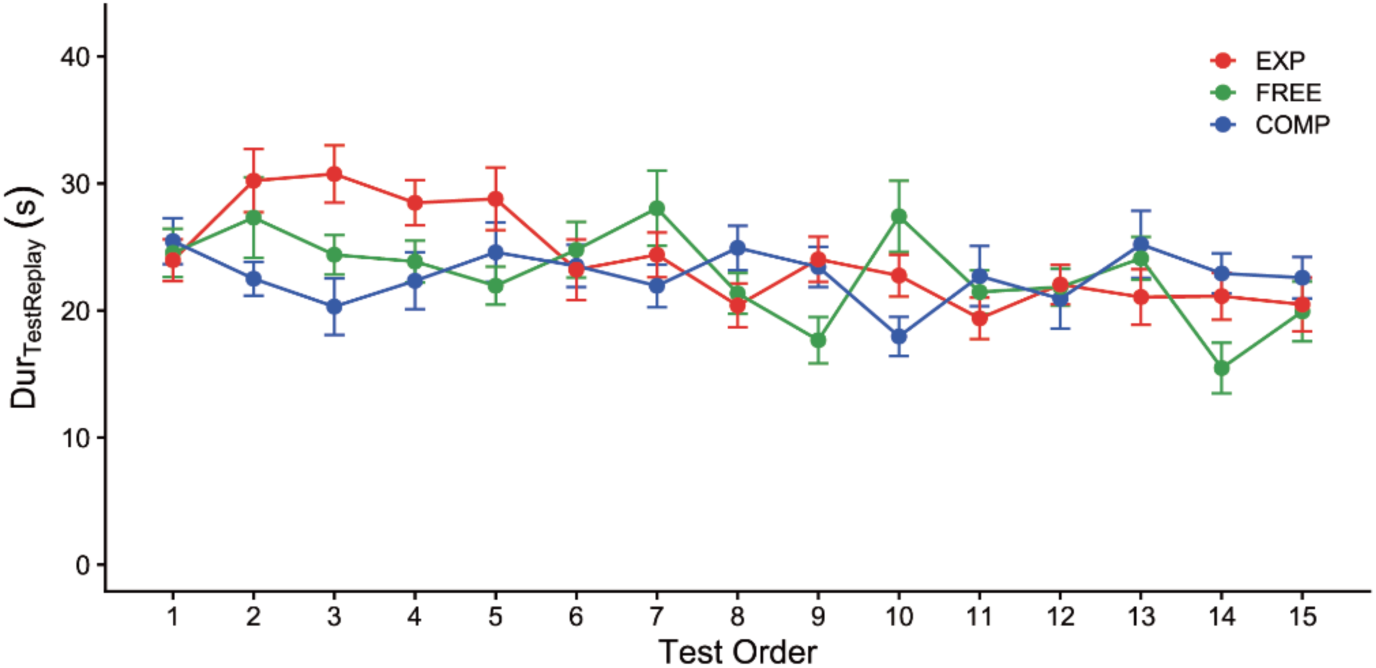
Duration_TestReplay_ during test phase for each subevent position (1-15), separated by prior manipulation condition and averaged across participants. Points show group means; error bars show ±1 SE. Compress: blue, Free: green, Expand: red.

**Supplementary Figure 6.**
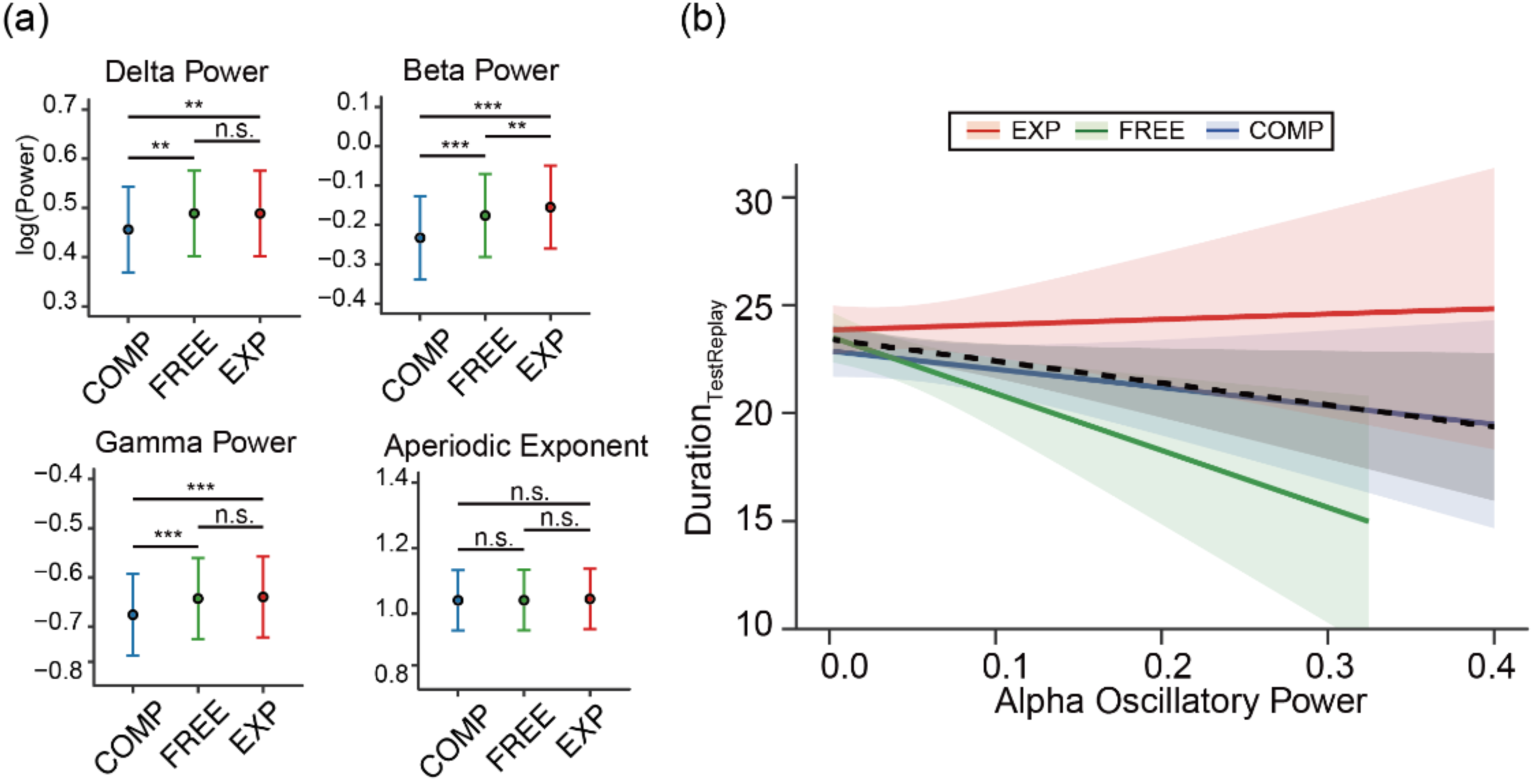
Additional oscillatory metrics and spontaneous replay duration. Similar to Fig. 3, (a) displays the estimated marginal means of log-transformed power across conditions for the remaining frequency bands (delta, beta, gamma) and the aperiodic exponent. Error bars indicate 95% CI. Omnibus tests revealed significant oscillatory power differences between manipulation conditions in the delta (*χ²*(2) = 14.382, *p* < .001), beta (*χ²*(2) = 131.305, *p* < .001), and gamma (*χ²*(2) = 41.559, *p* < .001) bands. Delta power was significantly lower in Compress than in Free (*b* = −0.030, *t* = −3.007, *p* = .008) and Expand (*b* = −0.034, *t* = −3.518, *p* = .001), with no Free–Expand difference (*p* = .875). Beta power increased monotonically from Compress to Expand, differing across all three pairs (Compress–Free: *b* = −0.056, *t* = −7.801, *p* < .001; Compress–Expand: *b* = −0.079, *t* = −11.190, *p* < .001; Free–Expand: *b* = −0.024, *t* = −3.353, *p* = .002). Gamma power was lower in Compress compared to Free (*b* = −0.030, *t* = −5.085, *p* < .001) and Expand (*b* = −0.036, *t* = −5.997, *p* < .001), with no Free–Expand difference (*p* = .653). The aperiodic exponent did not differ across any condition pairs (all *ps* > .86). (b) Alpha power during the manipulation phase and the time taken to replay each subevent at test (Duration_TestReplay_). Higher alpha power predicted shorter test replay (*b* = −0.833, *z* = −3.037, *p* = .002). This relationship did not depend on condition (frequency × condition interactions: *χ²*(4) = 2.02, *p* = .731), and an alpha-only model gave the best fit (lowest AIC and BIC); adding theta power, the aperiodic exponent or interaction terms did not improve it. The black dashed line shows the overall fixed effect; coloured lines show condition-specific fits. For all panels, condition color coding: Compress (blue), Free (green), Expand (red). Shaded ribbons represent 95% CI. \**p* < .05, \*\**p* < .01, \*\*\**p* < .001, n.s. = not significant.

**Supplementary Table 1.**
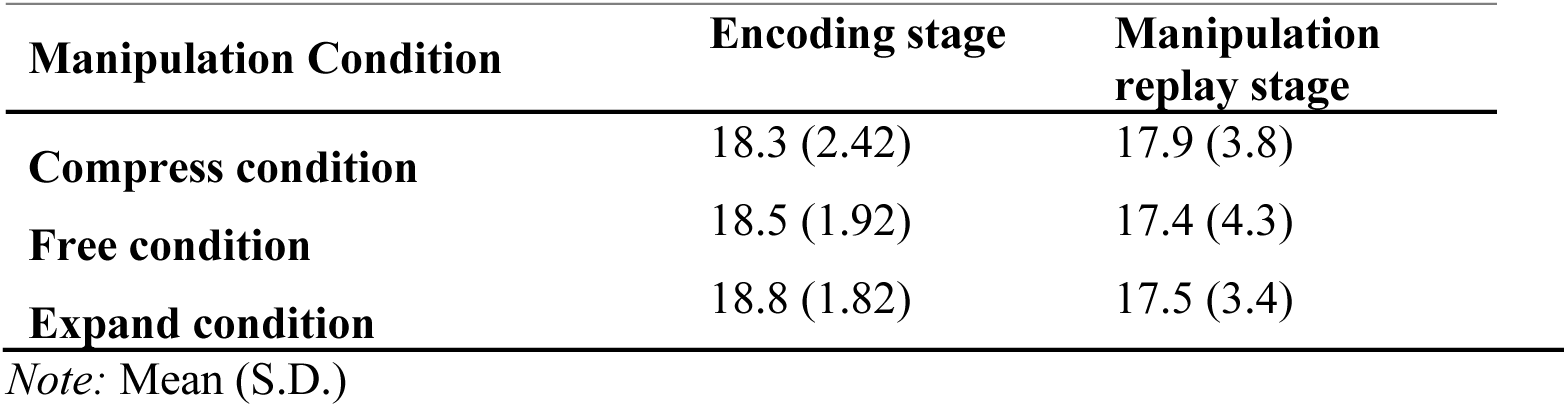
Number of Subevent Epochs after Filtering out of 20 Total Subevent.

**Supplementary Table 2.** GLMM Results for Duration_TestReplay_ (Effect of Condition) Formula: Duration_TestReplay_ ∼ Duration_ActualVideo_ + Condition + (1|Subject) + (1|Event)

| Predictor | Estimate (b) | SE | z-value | p-value | VIF |
| --- | --- | --- | --- | --- | --- |
| **(Intercept)** | 2.951 | 0.073 | 40.16 | < .001 | — |
| **Video Duration (50s vs 30s)** | 0.152 | 0.049 | 3.10 | < .001 | 1.000 |
| **Condition: Compress (vs Free)** | 0.014 | 0.033 | 0.41 | .679 | 1.000 |
| **Condition: Expand (vs Free)** | 0.046 | 0.033 | 1.38 | .166 | 1.000 |

Note. SE: Standard Error. VIF: Variance Inflation Factor. The model was fitted using a Gamma distribution with a log link function and includes random intercepts for Subject and Event. Reference levels: Condition = Free, Video Duration = 30s.

**Supplementary Table 3.** GLMM Results for Duration_TestReplay_ (Effect of Duration_Mani_) Formula: Duration_TestReplay_ ∼ Duration_ActualVideo_ + Duration_Mani_ (Z-scored) + (1|Subject) + (1|Event)

| Predictor | Estimate (b) | SE | z-value | P-value | VIF |
| --- | --- | --- | --- | --- | --- |
| **(Intercept)** | 2.989 | 0.067 | 44.36 | < .001 | — |
| **Video Duration (50s vs 30s)** | 0.118 | 0.049 | 2.41 | .016 | 1.15 |
| **Manipulation Duration (Z-scored)** | 0.074 | 0.017 | 4.45 | < .001 | 1.15 |

Note. SE: Standard Error. VIF: Variance Inflation Factor. The model was fitted using a Gamma distribution with a log link function and includes random intercepts for Subject and Event. Reference levels: Condition = Free, Video Duration = 30s.

**Supplementary Table 4.**
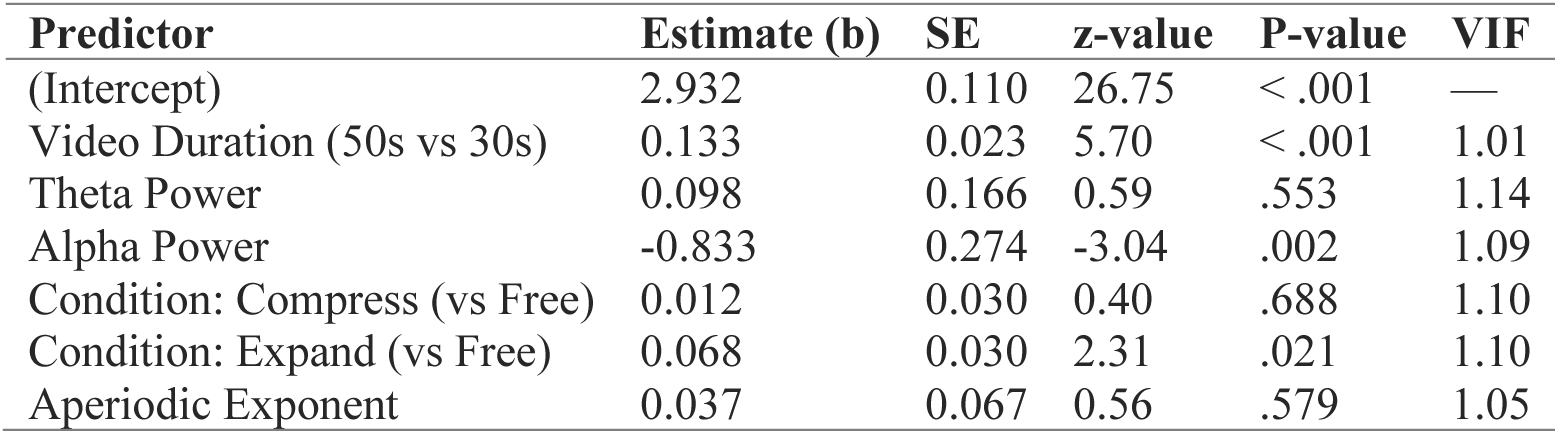
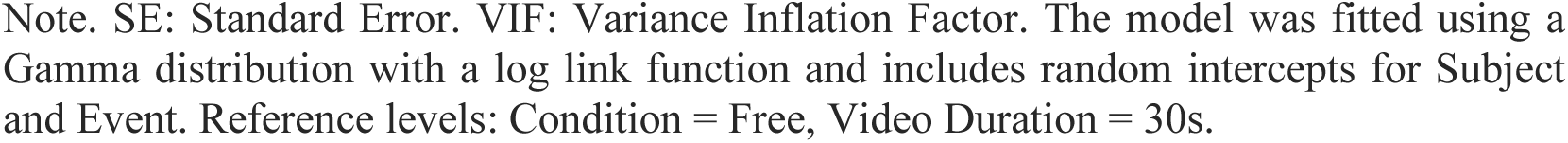
GLMM Results for Duration_TestReplay_ (Neural Oscillatory Predictors) Formula: Duration_TestReplay_ ∼ Duration_ActualVideo_ + Theta Oscillatory Power + Alpha Oscillatory Power + Condition + (1|Subject) + (1|Event)

| Predictor | Estimate (b) | SE | z-value | P-value | VIF |
| --- | --- | --- | --- | --- | --- |
| **(Intercept)** | 2.932 | 0.110 | 26.75 | < .001 | — |
| **Video Duration (50s vs 30s)** | 0.133 | 0.023 | 5.70 | < .001 | 1.01 |
| **Theta Power** | 0.098 | 0.166 | 0.59 | .553 | 1.14 |
| **Alpha Power** | -0.833 | 0.274 | -3.04 | .002 | 1.09 |
| **Condition: Compress (vs Free)** | 0.012 | 0.030 | 0.40 | .688 | 1.10 |
| **Condition: Expand (vs Free)** | 0.068 | 0.030 | 2.31 | .021 | 1.10 |
| **Aperiodic Exponent** | 0.037 | 0.067 | 0.56 | .579 | 1.05 |

Note. SE: Standard Error. VIF: Variance Inflation Factor. The model was fitted using a Gamma distribution with a log link function and includes random intercepts for Subject and Event. Reference levels: Condition = Free, Video Duration = 30s.

**Supplementary Table 5.**
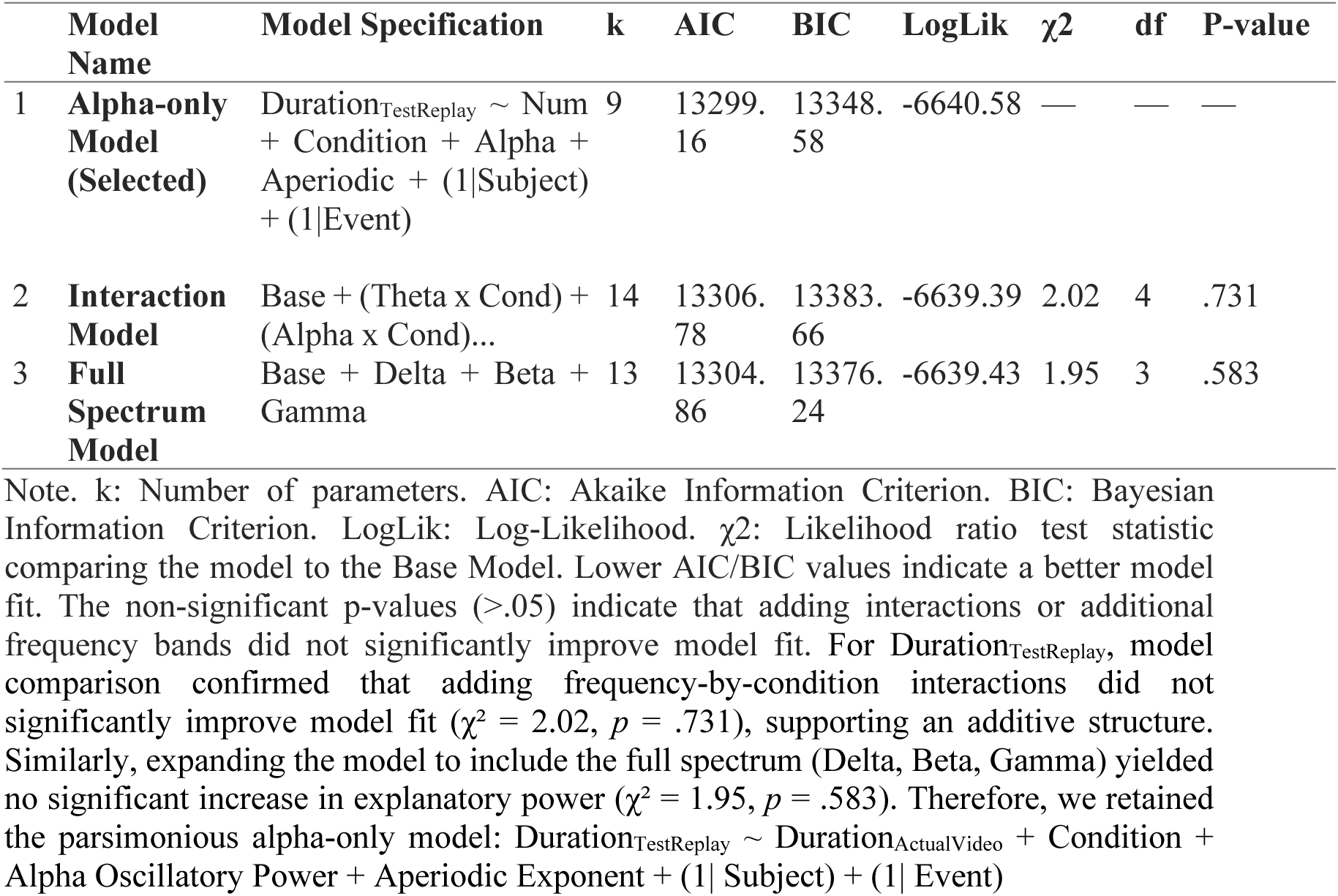
GLMM comparison for Duration_TestReplay_ (Neural Oscillatory Predictors)

**Supplementary Table 6.** GLMM Results for Duration_TestReported_ (Effect of Condition) Formula: Duration_TestReported_ ∼ Duration_ActualVideo_ + Condition + (1|Subject) + (1|Event)

| Predictor | Estimate (b) | SE | z-value | P-value | VIF |
| --- | --- | --- | --- | --- | --- |
| (Intercept) | 2.729 | 0.101 | 27.15 | < .001 | — |
| Video Duration (50s vs 30s) | 0.210 | 0.044 | 4.77 | < .001 | 1.000 |
| Condition: Compress (vs Free) | -0.026 | 0.019 | -1.33 | .182 | 1.000 |
| Condition: Expand (vs Free) | 0.055 | 0.020 | 2.79 | .005 | 1.000 |

Note. SE: Standard Error. VIF: Variance Inflation Factor. The model was fitted using a Gamma distribution with a log link function and includes random intercepts for Subject and Event. Reference levels: Condition = Free, Video Duration = 30s.

**Supplementary Table 7.** GLMM Results for Duration_TestReported_ (Effect of Duration_Mani_) Formula: Duration_TestReported_ ∼ Duration_ActualVideo_ + Duration_Mani_ (Z-scored) + (1|Subject) + (1|Event)

| Predictor | Estimate (b) | SE | z-value | P-value | VIF |
| --- | --- | --- | --- | --- | --- |
| (Intercept) | 2.758 | 0.097 | 28.56 | < .001 | — |
| Video Duration (50s vs 30s) | 0.174 | 0.042 | 4.13 | < .001 | 1.07 |
| Manipulation Duration (Z-scored) | 0.082 | 0.010 | 8.37 | < .001 | 1.07 |

Note. SE: Standard Error. VIF: Variance Inflation Factor. The model was fitted using a Gamma distribution with a log link function and includes random intercepts for Subject and Event. Reference levels: Condition = Free, Video Duration = 30s.

**Supplementary Table 8.** GLMM Results for Duration_TestReported_ (Neural Oscillatory Predictors × Condition) Formula: Duration_TestReported_ ∼ Theta Oscillatory Power * Condition + Alpha * Condition Oscillatory Power + (1|Subject) + (1|Event)

| Predictor | Estimate (b) | SE | z-value | P-value | GVIF |
| --- | --- | --- | --- | --- | --- |
| (Intercept) | 2.838 | 0.112 | 25.40 | < .001 | — |
| Video Duration (50s vs 30s) | 0.199 | 0.014 | 14.32 | < .001 | 1.01 |
| Condition Main Effects (Ref: Free) |  |  |  |  |  |
| Condition: Compress | -0.054 | 0.025 | -2.21 | .027 | 3.51 |
| Condition: Expand | 0.014 | 0.023 | 0.60 | .550 | 3.51 |
| Oscillatory Main Effects (in Free) |  |  |  |  |  |
| Theta Power | -0.057 | 0.158 | -0.36 | .718 | 2.88 |
| Alpha Power | -0.445 | 0.281 | -1.58 | .113 | 3.33 |
| Theta Interactions (Ref: Free) |  |  |  |  |  |
| Theta x Condition: Compress | -0.001 | 0.186 | -0.00 | .996 | 3.51 |
| Theta x Condition: Expand | 0.503 | 0.215 | 2.35 | .019 | 3.51 |
| Alpha Interactions (Ref: Free) |  |  |  |  |  |
| Alpha x Condition: Compress | 0.802 | 0.309 | 2.60 | .009 | 3.33 |
| Alpha x Condition: Expand | 0.078 | 0.335 | 0.23 | .816 | 3.33 |

Note. SE: Standard Error. GVIF: Generalized Variance Inflation Factor. All values were < 5, indicating no problematic multicollinearity. The model was fitted using a Gamma distribution with a log link function and includes random intercepts for Subject and Event. Reference levels: Condition = Free, Video Duration = 30s.

**Supplementary Table 9.**
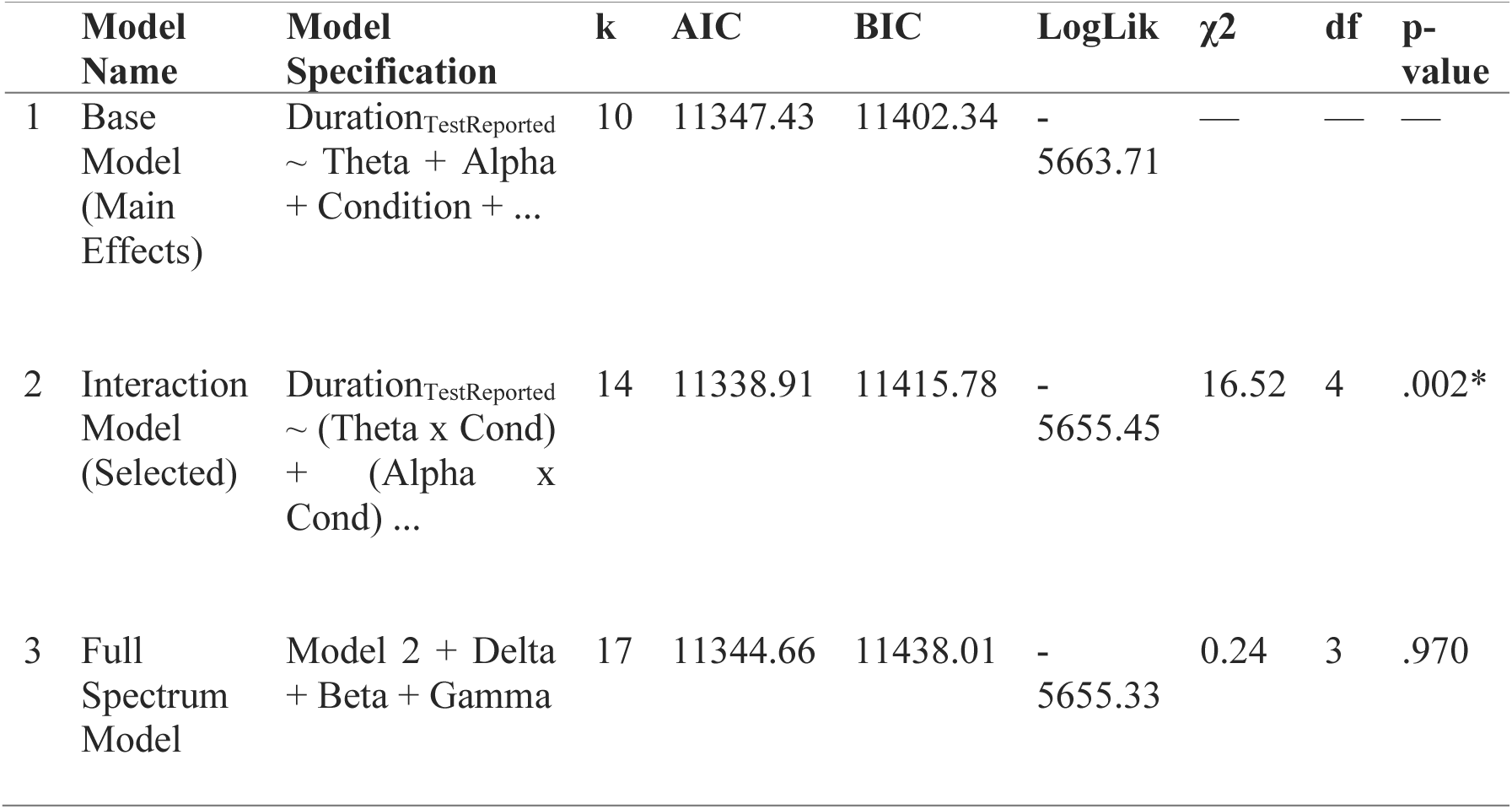
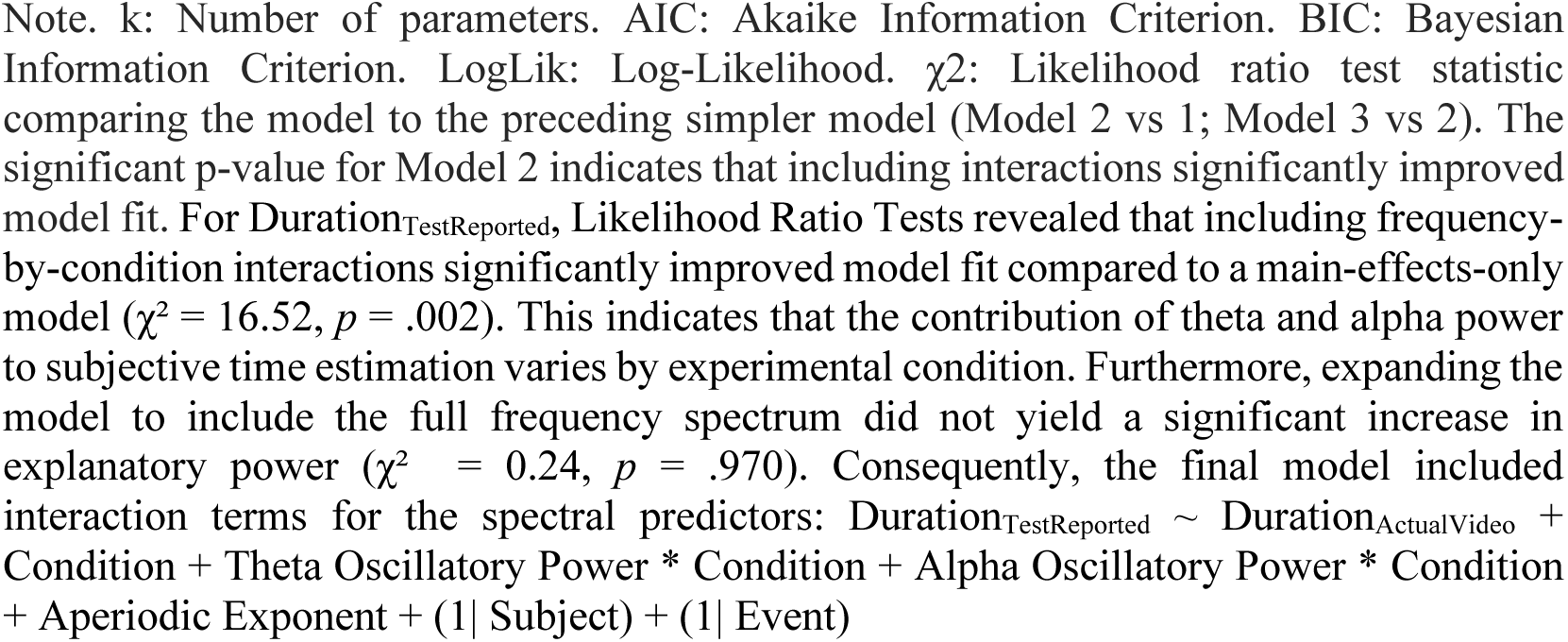
GLMM comparison for Duration_TestReported_ (Neural Oscillatory Predictors)

|  | Model Name | Model Specification | k | AIC | BIC | LogLik | χ<sup>2</sup> | df | p-value |
| --- | --- | --- | --- | --- | --- | --- | --- | --- | --- |
| 1 | Base Model (Main Effects) | Duration<sub>TestReported</sub> ~ Theta + Alpha + Condition + ... | 10 | 11347.43 | 11402.34 | -5663.71 | — | — | — |
| 2 | Interaction Model (Selected) | Duration<sub>TestReported</sub> ~ (Theta x Cond) + (Alpha x Cond) ... | 14 | 11338.91 | 11415.78 | -5655.45 | 16.52 | 4 | .002* |
| 3 | Full Spectrum Model | Model 2 + Delta + Beta + Gamma | 17 | 11344.66 | 11438.01 | -5655.33 | 0.24 | 3 | .970 |

Note. k: Number of parameters. AIC: Akaike Information Criterion. BIC: Bayesian Information Criterion. LogLik: Log-Likelihood. $\chi^2$ : Likelihood ratio test statistic comparing the model to the preceding simpler model (Model 2 vs 1; Model 3 vs 2). The significant p-value for Model 2 indicates that including interactions significantly improved model fit. For $\text{Duration}_{\text{TestReported}}$ , Likelihood Ratio Tests revealed that including frequency-by-condition interactions significantly improved model fit compared to a main-effects-only model ( $\chi^2 = 16.52, p = .002$ ). This indicates that the contribution of theta and alpha power to subjective time estimation varies by experimental condition. Furthermore, expanding the model to include the full frequency spectrum did not yield a significant increase in explanatory power ( $\chi^2 = 0.24, p = .970$ ). Consequently, the final model included interaction terms for the spectral predictors: $\text{Duration}_{\text{TestReported}} \sim \text{Duration}_{\text{ActualVideo}} + \text{Condition} + \text{Theta Oscillatory Power} * \text{Condition} + \text{Alpha Oscillatory Power} * \text{Condition} + \text{Aperiodic Exponent} + (1 | \text{Subject}) + (1 | \text{Event})$

**Supplementary Table 10.** GLMM Comparison Testing the Independence of Oscillatory Effects from Memory Content.

| | Model Name | Model Specification | k | AIC | BIC | LogLik | $\chi^2$ | p-value | BF01 |
| --- | --- | --- | --- | --- | --- | --- | --- | --- | --- |
| 1 | Interaction Model | $\text{Duration}_{\text{TestReported}} \sim (\text{Theta} \times \text{Cond}) + (\text{Alpha} \times \text{Cond}) \dots$ | 14 | 11338.91 | 11415.78 | -5655.45 | — | — | — |
| 2 | Independence Model (+MTS) | Model 1 + MTS | 15 | 11337.80 | 11420.17 | -5653.90 | 3.11 | .078 | 8.96 |

Note. k: Number of parameters. AIC: Akaike Information Criterion. BIC: Bayesian Information Criterion (lower is better). LogLik: Log-Likelihood. $\text{BF}_{01}$ : Bayes Factor quantifying evidence for the Null Hypothesis (Model 1) over the Alternative (Model 2). A $\text{BF}_{01}$ of 8.96 indicates moderate evidence that adding MTS does not improve the model.

**Supplementary Table 11.**
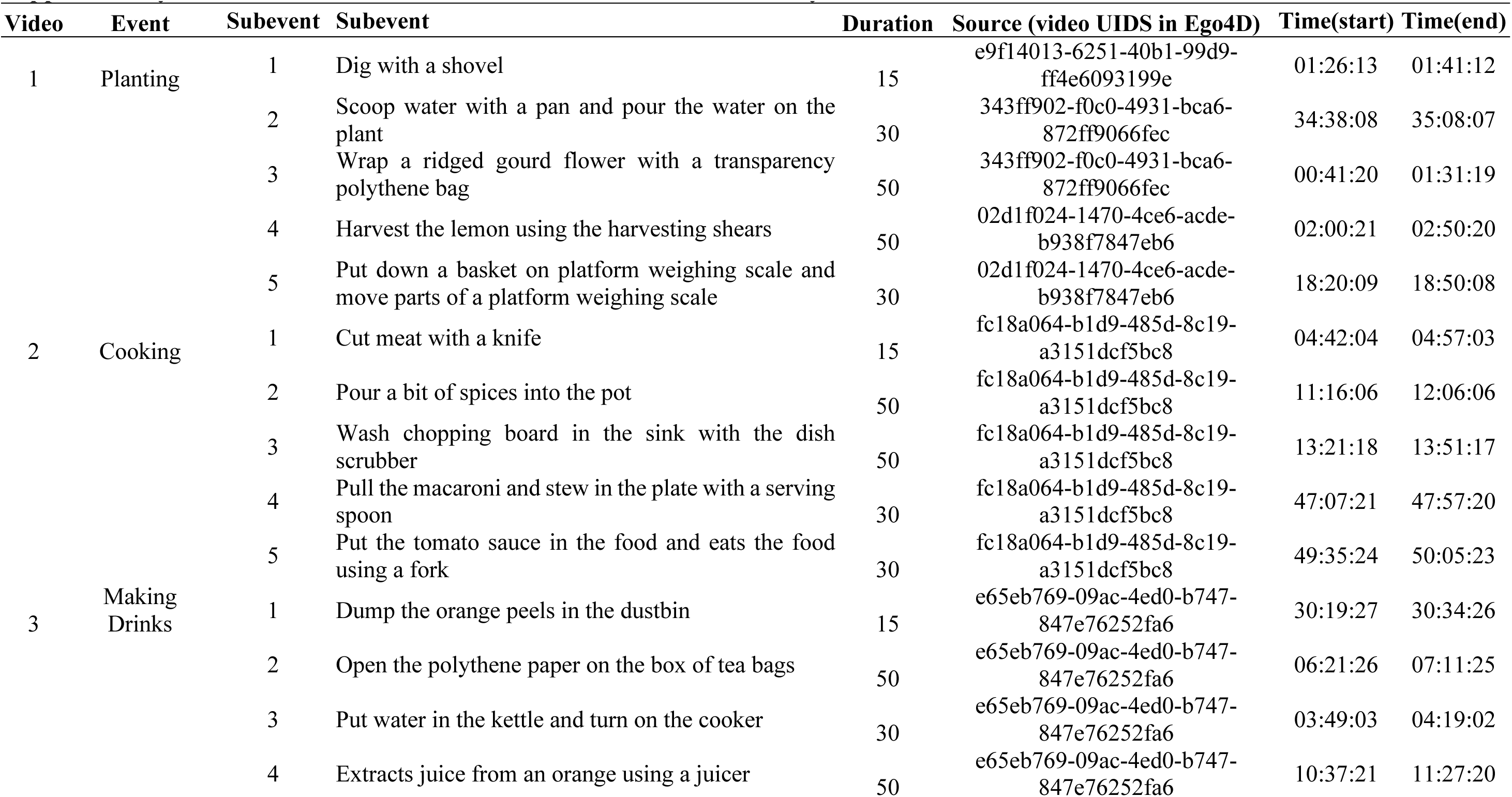
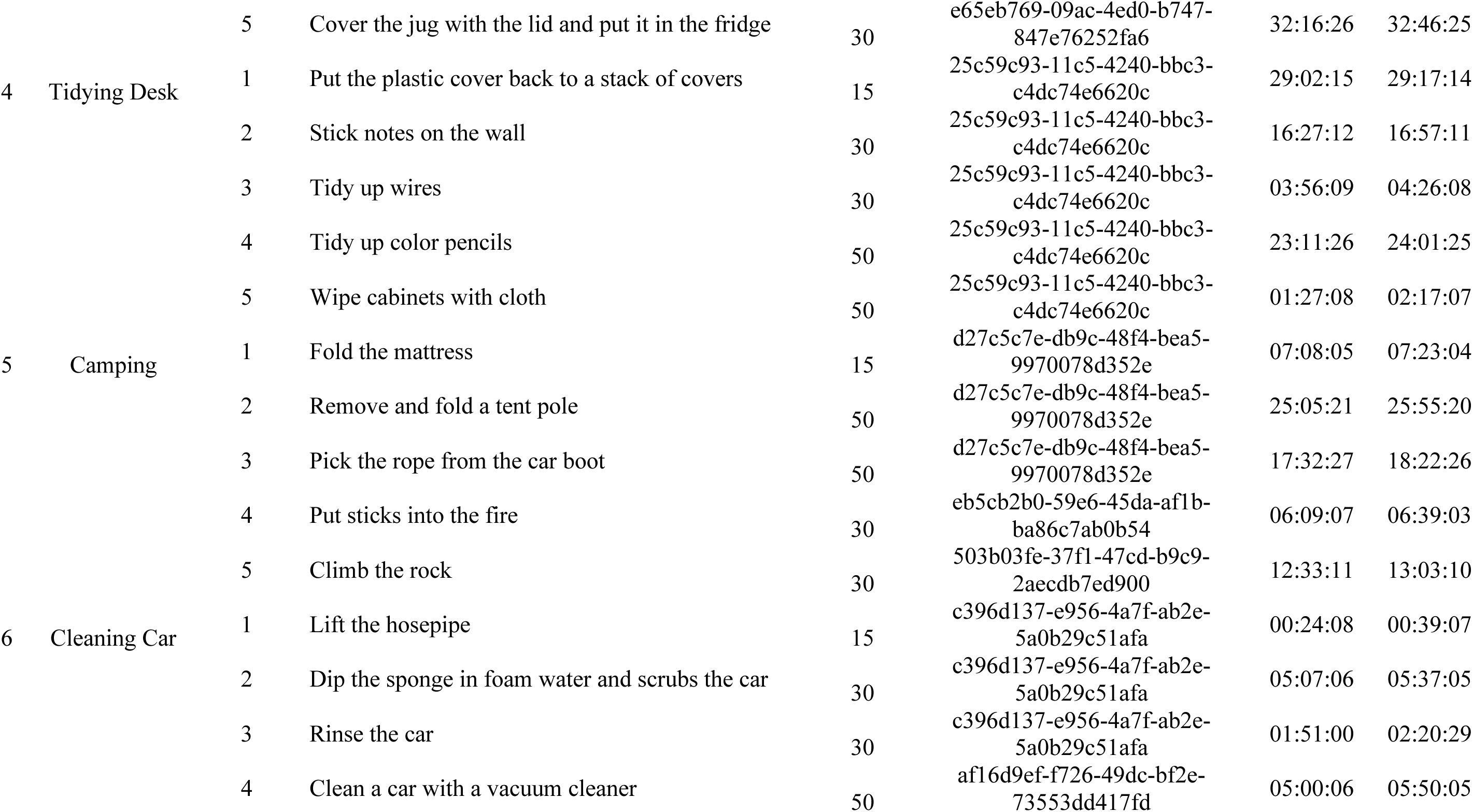
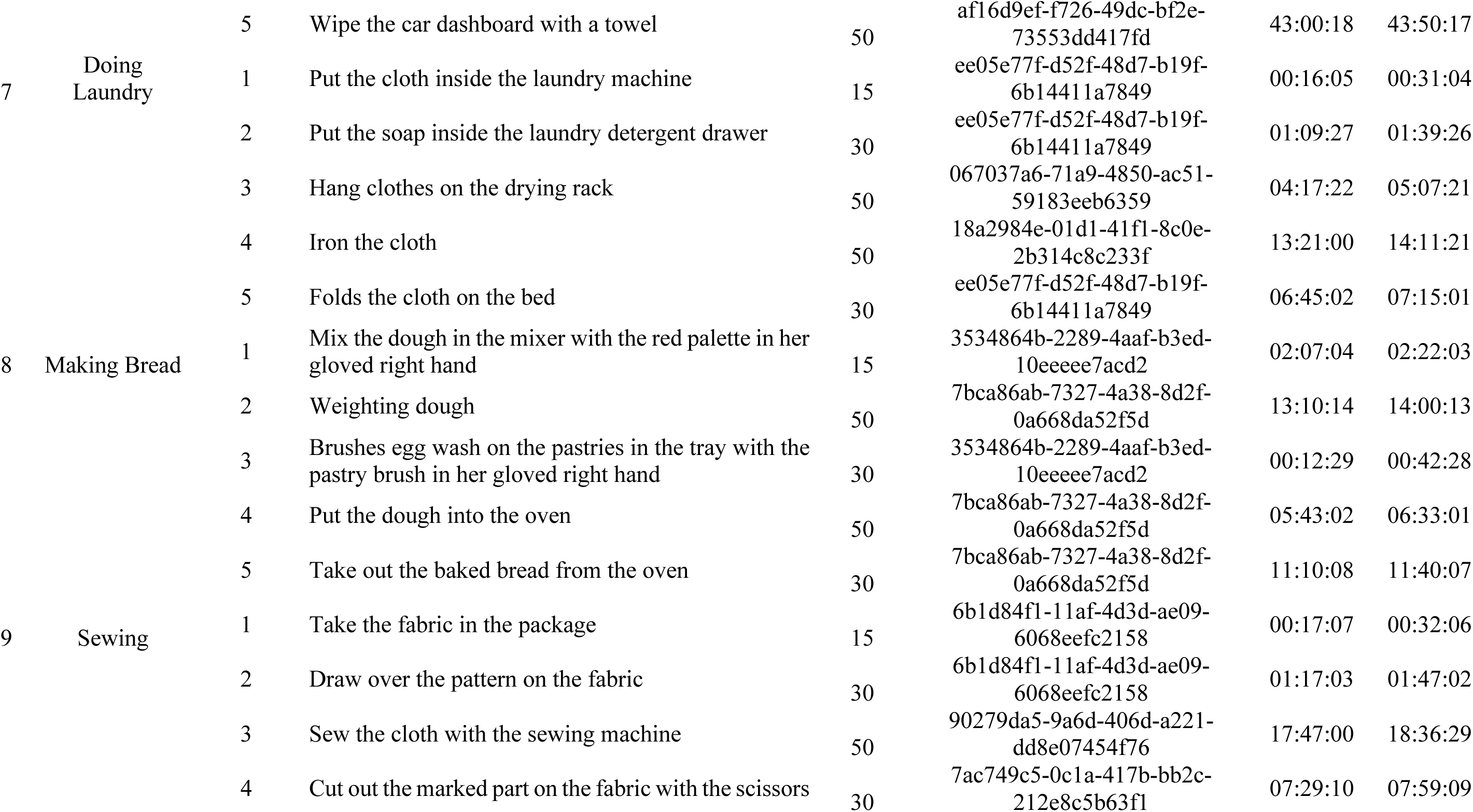
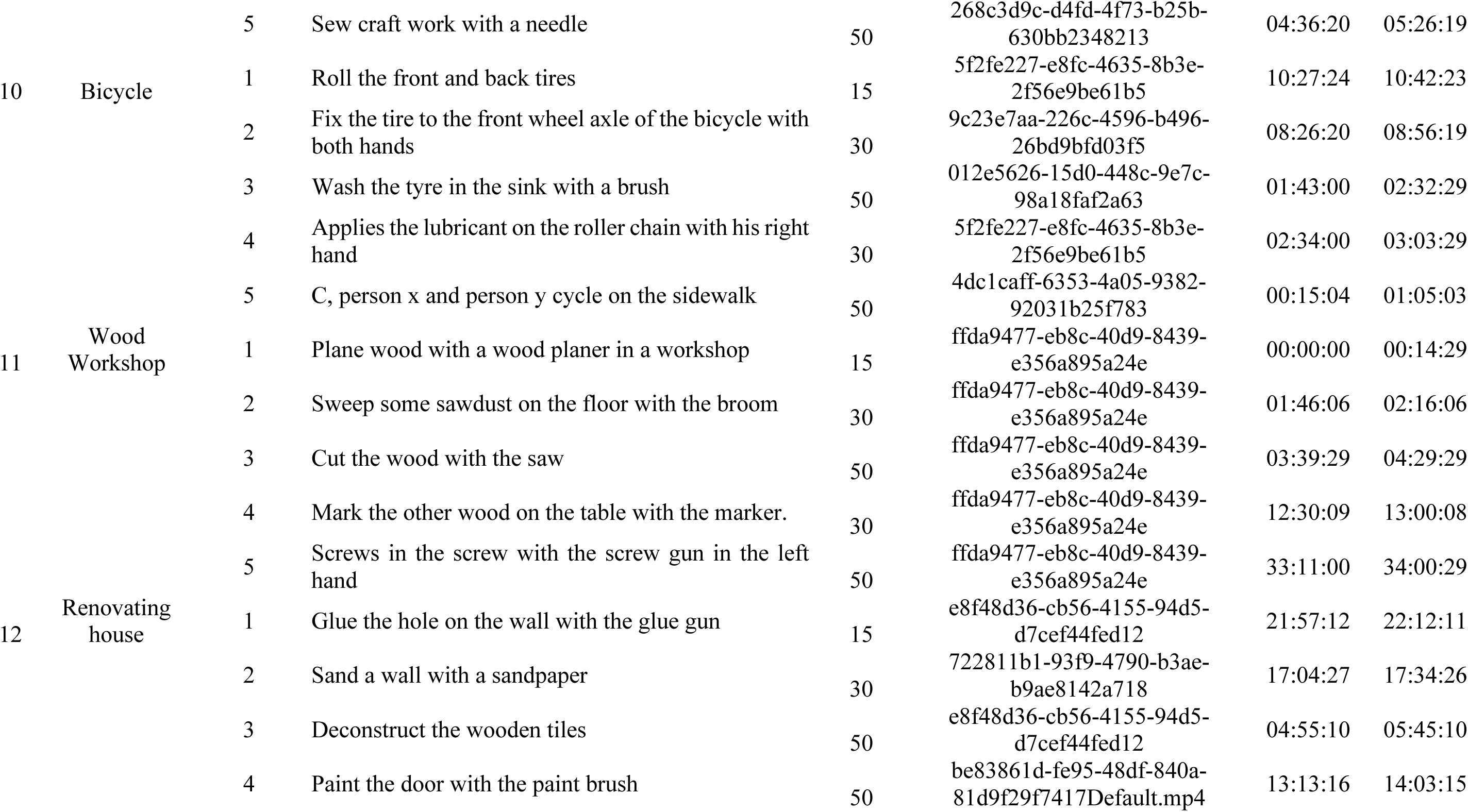
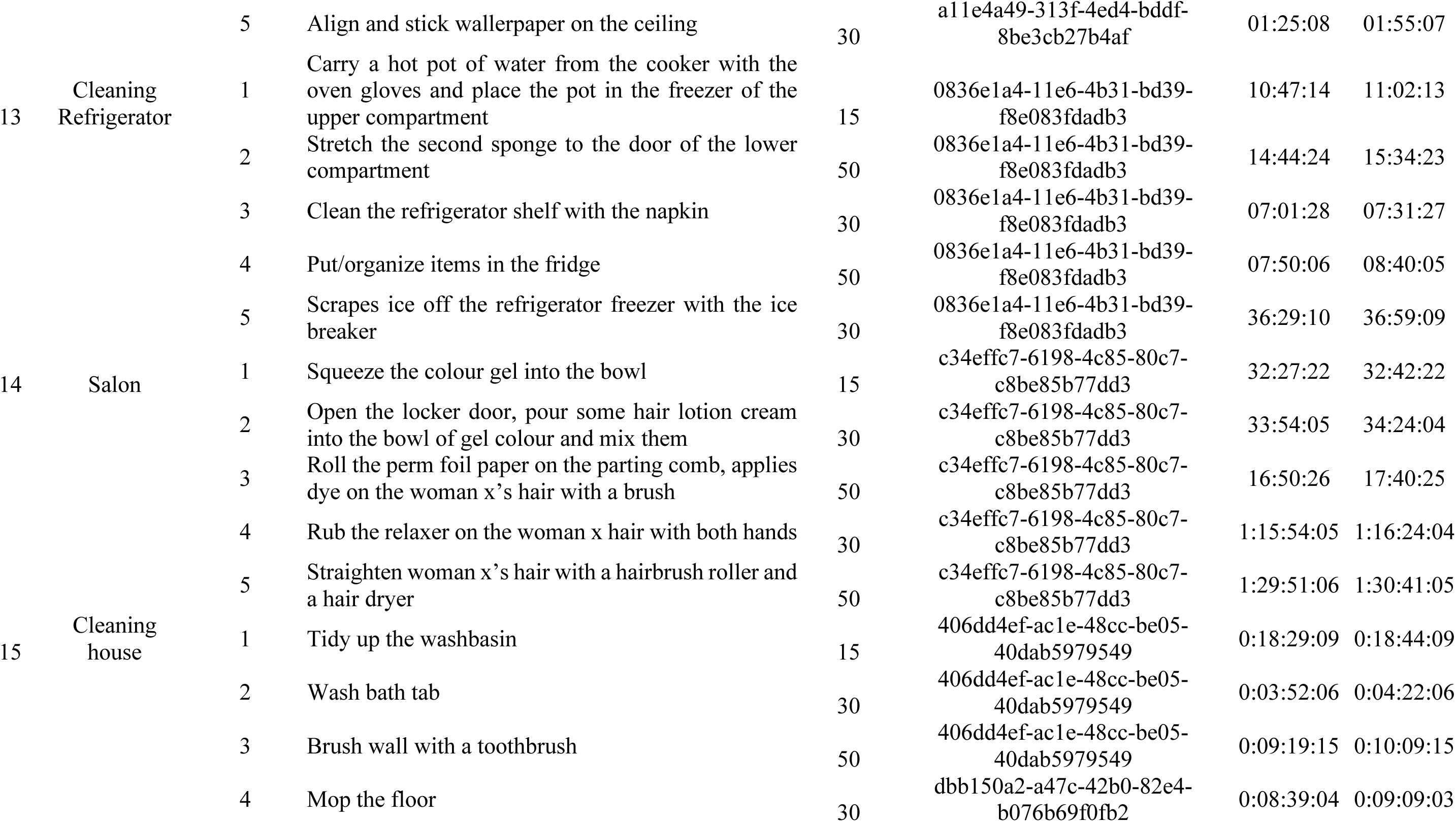
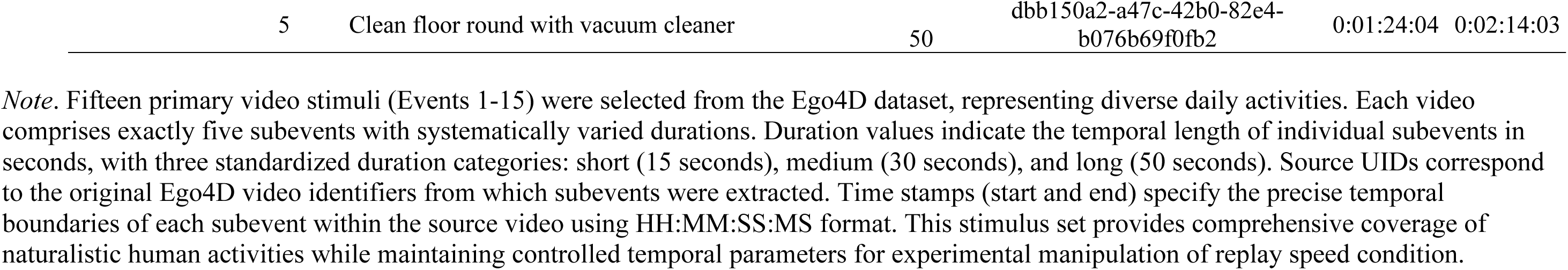
Characteristics of video stimuli used in the study.

## References

1. Azizi, L., Polti, I., & Wassenhove, V. van. (2023). Spontaneous α Brain Dynamics Track the Episodic “When”. Journal of Neuroscience, 43(43), 7186–7197.

2. Bastien, C. H., Vallières, A., & Morin, C. M. (2001). Validation of the Insomnia Severity Index as an outcome measure for insomnia research. Sleep medicine, 2(4), 297–307.

3. Bates, D., Mächler, M., Bolker, B., & Walker, S. (2015). Fitting linear mixed-effects models using lme4. Journal of Statistical Software, 67(1), 1–48

4. Bellingrath, J. E. (2023). The Self-Simulational Theory of temporal extension. Neuroscience of Consciousness, 2023(1), niad015.

5. Bellmund, J. L. S., Gärdenfors, P., Moser, E. I., & Doeller, C. F. (2018). Navigating cognition: Spatial codes for human thinking. Science, 362(6415), eaat6766.

6. Biderman, N., Bakkour, A., & Shohamy, D. (2020). What Are Memories For? The Hippocampus Bridges Past Experience with Future Decisions. Trends in Cognitive Sciences, 24(7), 542–556.

7. Block, R. A. (1992). Prospective and retrospective duration judgment: The role of information processing and memory. In F. Macar, V. Pouthas, & W. J. Friedman (Eds.), Time, Action and Cognition: Towards Bridging the Gap (pp. 141–152). Springer Netherlands.

8. Block, R. A., & Reed, M. A. (1978). Remembered duration: Evidence for a contextual-change hypothesis. Journal of Experimental Psychology: Human Learning and Memory, 4(6), 656–665.

9. Block, R. A., & Zakay, D. (1997). Prospective and retrospective duration judgments: A meta-analytic review. Psychonomic Bulletin & Review, 4(2), 184–197.

10. Brady, T. F., Konkle, T., Alvarez, G. A., & Oliva, A. (2013). Real-world objects are not represented as bound units: Independent forgetting of different object details from visual memory. Journal of Experimental Psychology: General, 142(3), 791–808.

11. Burt, C. D. B., & Kemp, S. (1991). Retrospective duration estimation of public events. Memory & Cognition, 19(3), 252–262.

12. Buysse DJ, Reynolds CF 3rd, Monk TH, Berman SR, Kupfer DJ. The Pittsburgh Sleep Quality Index: a new instrument for psychiatric practice and research. Psychiatry Res. 1989 May;28(2):193–213.

13. Buzsáki, G., & Moser, E. I. (2013). Memory, navigation and theta rhythm in the hippocampal-entorhinal system. Nature Neuroscience, 16(2), 130–138.

14. Buzsáki, G., & Tingley, D. (2018). Space and Time: The Hippocampus as a Sequence Generator. *Trends in Cognitive Sciences*, Special Issue: Time in the Brain, 22(10), 853–869.

15. Cao, R., Bladon, J. H., Charczynski, S. J., Hasselmo, M. E., & Howard, M. W. (2022). Internally generated time in the rodent hippocampus is logarithmically compressed. eLife, 11, e75353.

16. Clewett, D., DuBrow, S., & Davachi, L. (2019). Transcending time in the brain: How event memories are constructed from experience. Hippocampus, 29(3), 162–183.

17. D’Argembeau, A., Jeunehomme, O., & Stawarczyk, D. (2022). Slices of the past: How events are temporally compressed in episodic memory. Memory, 30(1), 43–48.

18. Delorme, A., & Makeig, S. (2004). EEGLAB: An open source toolbox for analysis of single-trial EEG dynamics including independent component analysis. Journal of Neuroscience Methods, 134(1), 9–21.

19. Denovellis, E. L., Gillespie, A. K., Coulter, M. E., Sosa, M., Chung, J. E., Eden, U. T., & Frank, L. M. (2021). Hippocampal replay of experience at real-world speeds. eLife, 10, e64505.

20. Donoghue, T., Haller, M., Peterson, E. J., Varma, P., Sebastian, P., Gao, R., Noto, T., Lara, A. H., Wallis, J. D., Knight, R. T., Shestyuk, A., & Voytek, B. (2020). Parameterizing neural power spectra into periodic and aperiodic components. Nature Neuroscience, 23(12), 1655–1665.

21. Eagleman, D. M., & Pariyadath, V. (2009). Is subjective duration a signature of coding efficiency? Philosophical Transactions of the Royal Society B: Biological Sciences, 364(1525), 1841–1851.

22. Eichenbaum, H. (2014). Time cells in the hippocampus: A new dimension for mapping memories. Nature Reviews Neuroscience, 15(11), 732–744.

23. El Haj, M., & Kapogiannis, D. (2016a). Time distortions in Alzheimer’s disease: A systematic review and theoretical integration. Npj Aging and Mechanisms of Disease, 2(1), 1–5.

24. El Haj, M., & Kapogiannis, D. (2016b). Time distortions in Alzheimer’s disease: A systematic review and theoretical integration. NPJ Aging and Mechanisms of Disease, 2, 16016.

25. El Haj, M., Moroni, C., Samson, S., Fasotti, L., & Allain, P. (2013). Prospective and retrospective time perception are related to mental time travel: Evidence from Alzheimer’s disease. Brain and Cognition, 83(1), 45–51.

26. Ezzyat, Y., & Davachi, L. (2014). Similarity Breeds Proximity: Pattern Similarity within and across Contexts Is Related to Later Mnemonic Judgments of Temporal Proximity. Neuron, 81(5), 1179–1189.

27. Faber, M., & Gennari, S. P. (2015). In search of lost time: Reconstructing the unfolding of events from memory. Cognition, 143, 193–202.

28. Fox, J., & Weisberg, S. (2019). An R companion to applied regression (3rd ed.). Sage.

29. Fuentemilla, L., Barnes, G. R., Düzel, E., & Levine, B. (2014). Theta oscillations orchestrate medial temporal lobe and neocortex in remembering autobiographical memories. NeuroImage, 85 Pt 2, 730–737.

30. Gao, R., Peterson, E. J., & Voytek, B. (2017). Inferring synaptic excitation/inhibition balance from field potentials. NeuroImage, 158, 70–78.

31. Grauman, K., Westbury, A., Byrne, E., Chavis, Z., Furnari, A., Girdhar, R., Hamburger, J., Jiang, H., Liu, M., Liu, X., Martin, M., Nagarajan, T., Radosavovic, I., Ramakrishnan, S. K., Ryan, F., Sharma, J., Wray, M., Xu, M., Xu, E. Z., … Malik, J. (2022). Ego4D: Around the World in 3,000 Hours of Egocentric Video (arXiv:2110.07058). arXiv.

32. Griffiths, B. J., Mayhew, S. D., Mullinger, K. J., Jorge, J., Charest, I., Wimber, M., & Hanslmayr, S. (2019). Alpha/beta power decreases track the fidelity of stimulus-specific information. eLife, 8, e49562.

33. Hanslmayr, S., Staudigl, T., & Fellner, M.-C. (2012). Oscillatory power decreases and long-term memory: The information via desynchronization hypothesis. Frontiers in Human Neuroscience, 6.

34. Herweg, N. A., Solomon, E. A., & Kahana, M. J. (2020). Theta oscillations in human memory. Trends in Cognitive Sciences, 24(3), 208–227.

35. Heusser, A. C., Poeppel, D., Ezzyat, Y., & Davachi, L. (2016). Episodic sequence memory is supported by a theta-gamma phase code. Nature Neuroscience, 19(10), 1374–1380.

36. Howard, M. W. (2018). Memory as perception of the past: Compressed time in mind and brain. Trends in Cognitive Sciences, 22(2), 124–136.

37. Hsieh, L.-T., Gruber, M. J., Jenkins, L. J., & Ranganath, C. (2014). Hippocampal Activity Patterns Carry Information about Objects in Temporal Context. Neuron, 81(5), 1165–1178.

38. Hupbach, A., Hardt, O., Gomez, R., & Nadel, L. (2008). The dynamics of memory: Context-dependent updating. Learning & Memory, 15(8), 574–579.

39. Jensen, O., Gelfand, J., Kounios, J., & Lisman, J. E. (2002). Oscillations in the alpha band (9-12 Hz) increase with memory load during retention in a short-term memory task. Cerebral Cortex, 12(8), 877–882.

40. Jensen, O., & Mazaheri, A. (2010). Shaping Functional Architecture by Oscillatory Alpha Activity: Gating by Inhibition. Frontiers in Human Neuroscience, 4, 186.

41. Jeunehomme, O., & D’Argembeau, A. (2019). The time to remember: Temporal compression and duration judgements in memory for real-life events. Quarterly Journal of Experimental Psychology, 72(4), 930–942.

42. Jeunehomme, O., Folville, A., Stawarczyk, D., Van Der Linden, M., & D’Argembeau, A. (2018). Temporal compression in episodic memory for real-life events. Memory, 26(6), 759–770.

43. Jeunehomme, O., Leroy, N., & D’Argembeau, A. (2020). The temporal compression of events during episodic future thinking. Cognition, 205, 104416.

44. Klimesch, W. (2012). Alpha-band oscillations, attention, and controlled access to stored information. Trends in Cognitive Sciences, 16(12), 606–617.

45. Klimesch, W., Sauseng, P., & Hanslmayr, S. (2007). EEG alpha oscillations: The inhibition-timing hypothesis. Brain Research Reviews, 53(1), 63–88.

46. Kononowicz, T. W., & van Wassenhove, V. (2016). In Search of Oscillatory Traces of the Internal Clock. Frontiers in Psychology, 7, 224.

47. Kuznetsova, A., Brockhoff, P. B., & Christensen, R. H. B. (2017). lmerTest package: Tests in linear mixed effects models. Journal of Statistical Software, 82(13), 1–26.

48. Lenth, R. V. (2023). *emmeans: Estimated marginal means, aka least-squares means* (Version 1.11.0) [Computer software]. https://CRAN.R-project.org/package=emmeans

49. Leroy, N., Majerus, S., & D’Argembeau, A. (2024). Working memory capacity for continuous events: The root of temporal compression in episodic memory? Cognition, 247, 105789.

50. Lisman, J. E., & Jensen, O. (2013). The θ-γ neural code. Neuron, 77(6), 1002–1016.

51. Liu, J., Zhang, H., Yu, T., Ren, L., Ni, D., Yang, Q., Lu, B., Zhang, L., Axmacher, N., & Xue, G. (2021). Transformative neural representations support long-term episodic memory. Science Advances, 7(41), eabg9715.

52. Lositsky, O., Chen, J., Toker, D., Honey, C. J., Shvartsman, M., Poppenk, J. L., Hasson, U., & Norman, K. A. (2016). Neural pattern change during encoding of a narrative predicts retrospective duration estimates. eLife, 5, e16070.

53. Lovibond, P. F., & Lovibond, S. H. (1995). The structure of negative emotional states: Comparison of the Depression Anxiety Stress Scales (DASS) with the Beck Depression and Anxiety Inventories. Behaviour Research and Therapy, 33(3), 335–343.

54. MacDonald, C. J., Lepage, K. Q., Eden, U. T., & Eichenbaum, H. (2011). Hippocampal “time cells” bridge the gap in memory for discontiguous events. Neuron, 71(4), 737–749.

55. Manza, P., Hau, C. L. V., & Leung, H.-C. (2014). Alpha Power Gates Relevant Information during Working Memory Updating. Journal of Neuroscience, 34(17), 5998–6002.

56. Matthews, W. J., & Meck, W. H. (2016). Temporal cognition: Connecting subjective time to perception, attention, and memory. Psychological Bulletin, 142(8), 865–907.

57. Mau, W., Sullivan, D. W., Kinsky, N. R., Hasselmo, M. E., Howard, M. W., & Eichenbaum, H. (2018). The same hippocampal CA1 population simultaneously codes temporal information over multiple timescales. Current Biology, 28(10), 1499–1508.e4.

58. Michelmann, S., Staresina, B. P., Bowman, H., & Hanslmayr, S. (2019). Speed of time-compressed forward replay flexibly changes in human episodic memory. Nature Human Behaviour, 3(2), 143–154.

59. Mioni, G., Grondin, S., & Stablum, F. (2014). Temporal dysfunction in traumatic brain injury patients: Primary or secondary impairment? Frontiers in Human Neuroscience, 8, 269.

60. Nadel, L., Samsonovich, A., Ryan, L., & Moscovitch, M. (2000). Multiple trace theory of human memory: Computational, neuroimaging, and neuropsychological results. Hippocampus, 10(4), 352–368.

61. Ornstein, R. E. (1969). On the Experience of Time. Harmondsworth.

62. Pastalkova, E., Itskov, V., Amarasingham, A., & Buzsáki, G. (2008). Internally Generated Cell Assembly Sequences in the Rat Hippocampus. Science, 321(5894), 1322–1327.

63. Pion-Tonachini, L., Kreutz-Delgado, K., & Makeig, S. (2019). ICLabel: An automated electroencephalographic independent component classifier, dataset, and website. NeuroImage, 198, 181–197.

64. Poynter, W. D. (1983). Duration judgment and the segmentation of experience. Memory & Cognition, 11(1), 77–82.

65. R Core Team. (2023). R: A language and environment for statistical computing (Version 4.3.0) [Computer software]. R Foundation for Statistical Computing. https://www.R-project.org/

66. Reddy, L., Zoefel, B., Possel, J. K., Peters, J., Dijksterhuis, D. E., Poncet, M., van Straaten, E. C. W., Baayen, J. C., Idema, S., & Self, M. W. (2021). Human hippocampal neurons track moments in a sequence of events. The Journal of Neuroscience, 41(31), 6714–6725.

67. Richards, B. A., & Frankland, P. W. (2017). The Persistence and Transience of Memory. Neuron, 94(6), 1071–1084.

68. Ritchey, M., Wing, E. A., LaBar, K. S., & Cabeza, R. (2013). Neural Similarity Between Encoding and Retrieval is Related to Memory Via Hippocampal Interactions. Cerebral Cortex, 23(12), 2818–2828.

69. Schacter, D. L., Addis, D. R., & Buckner, R. L. (2007). Remembering the past to imagine the future: The prospective brain. Nature Reviews Neuroscience, 8(9), 657–661.

70. Seeber, M., Stangl, M., Vallejo Martelo, M., Topalovic, U., Hiller, S., Halpern, C. H., Langevin, J.-P., Rao, V. R., Fried, I., Eliashiv, D., & Suthana, N. (2025). Human neural dynamics of real-world and imagined navigation. Nature Human Behaviour, 9(4), 781–793.

71. Strauß, A., Wöstmann, M., & Obleser, J. (2014). Cortical alpha oscillations as a tool for auditory selective inhibition. Frontiers in Human Neuroscience, 8.

72. Tingley, D., Yamamoto, T., Hirose, K., Keele, L., & Imai, K. (2014). mediation: R package for causal mediation analysis. Journal of Statistical Software, 59(5), 1–38.

73. Tsao, A., Sugar, J., Lu, L., Wang, C., Knierim, J. J., Moser, M.-B., & Moser, E. I. (2018). Integrating time from experience in the lateral entorhinal cortex. Nature, 561(7721), 57–62.

74. Tu, C.-A., Parviainen, T., Hämäläinen, J. A., & Hsu, Y.-F. (2025). Alpha oscillations protect auditory working memory against distractors in the encoding phase. Neuropsychologia, 207, 109058.

75. Umbach, G., Kantak, P., Jacobs, J., Kahana, M., Pfeiffer, B. E., Sperling, M., & Lega, B. (2020). Time cells in the human hippocampus and entorhinal cortex support episodic memory. Proceedings of the National Academy of Sciences, 117(45), 28463–28474.

76. Vass, L. K., Copara, M. S., Seyal, M., Shahlaie, K., Farias, S. T., Shen, P. Y., & Ekstrom, A. D. (2016). Oscillations Go the Distance: Low-Frequency Human Hippocampal Oscillations Code Spatial Distance in the Absence of Sensory Cues during Teleportation. Neuron, 89(6), 1180–1186.

77. Vicario, C. M., & Felmingham, K. L. (2018). Slower Time estimation in Post-Traumatic Stress Disorder. Scientific Reports, 8(1), 392.

78. Whittington, J. C. R., Muller, T. H., Mark, S., Chen, G., Barry, C., Burgess, N., & Behrens, T. E. J. (2020). The Tolman-Eichenbaum Machine: Unifying Space and Relational Memory through Generalization in the Hippocampal Formation. Cell, 183(5), 1249–1263.e23.

79. Wickham, H. (2016). ggplot2: Elegant graphics for data analysis (2nd ed.). Springer.

80. Wickham, H., François, R., Henry, L., Müller, K., & Vaughan, D. (2023). *dplyr: A grammar of data manipulation* (Version 1.1.4) [Computer software]. https://CRAN.R-project.org/package=dplyr

81. Yonelinas, A. P. (2002). The Nature of Recollection and Familiarity: A Review of 30 Years of Research. Journal of Memory and Language, 46(3), 441–517.

82. Yue, W. W. Y., Liu, J., Yao, Z., Zhang, Y., Mu, Z., & Hu, X. (2026). Retrospective duration judgments of naturalistic events depend on memories of event boundaries. Psychonomic Bulletin & Review, 33(1), 33.

83. Zakay, D., & Block, R. A. (2004). Prospective and retrospective duration judgments: An executive-control perspective. Acta Neurobiologiae Experimentalis, 64(3), 319–328.

84. Zhu, Z., Wang, Y., Jia, J., & Wu, Y. (2019). Beta and alpha desynchronizations underlie reconsolidation-mediated episodic memory updating. Neuropsychologia, 132, 107135.

